# Design Principles for Inflammasome Inhibition by Pyrin-Only-Proteins

**DOI:** 10.1101/2022.08.02.502519

**Authors:** Zachary Mazanek, Shuai Wu, Gretchen Belotte, Jeffery J. Zhou, Christina M. Stallings, Archit Garg, Jacob Lueck, Jungsan Sohn

## Abstract

Inflammasomes are filamentous signaling platforms essential for host defense against various intracellular calamities such as pathogen invasion and genotoxic stresses. However, dysregulated inflammasomes cause an array of human diseases including autoinflammatory disorders and cancer. It was recently identified that endogenous pyrin-only-proteins (POPs) regulate inflammasomes by directly inhibiting their filament assembly. Here, by combining Rosetta *in silico*, *in vitro*, and *in cellulo* methods, we investigate the target specificity and inhibition mechanisms of POPs. In contrast to a previous report, we find that POP1 is a poor inhibitor of the central inflammasome adaptor ASC. Instead, POP1 inhibits the assembly of upstream receptor PYD filaments such as those of AIM2, IFI16, NLRP3, and NLRP6. Moreover, not only does POP2 directly suppress the nucleation of ASC, but it can also inhibit the elongation of receptor filaments. In addition to inhibiting the elongation of AIM2 and NLRP6 filaments, POP3 potently suppresses the nucleation of ASC. Our Rosetta analyses and biochemical experiments consistently suggest that a combination of favorable and unfavorable interactions between POPs and PYDs is necessary for effective recognition and inhibition. Together, we reveal the intrinsic target redundancy of POPs and their inhibitory mechanisms.

## Introduction

Inflammasomes are filamentous signaling platforms integral to host innate defense against a wide range of intracellular catastrophes, which include ionizing irradiation, genotoxic chemicals, and pathogen invasion (Broz & Dixit, 2016; Zheng et al., 2020). However, persisting inflammasome activities lead to several human maladies including numerous autoinflammatory disorders, cancer, and even severe COVID-19 (Karki et al., 2017; Tartey & Kanneganti, 2020; Vora et al., 2021). Thus, understanding how inflammasome assemblies are regulated at the molecular level can provide key insights into developing strategies for preventing and treating various diseases (Broz & Dixit, 2016; Karki et al., 2017; Tartey & Kanneganti, 2020; Vora et al., 2021; Zheng et al., 2020).

Inflammasomes transduce signals by sequentially assembling filamentous oligomers, with multiple initial pathways progressively converging at downstream assemblies (Broz & Dixit, 2016; Kagan et al., 2014; Lu & Wu, 2015; Zheng et al., 2020). For instance, an array of molecular signatures arising from various pathogenic conditions induces the oligomerization of inflammasome receptors, resulting in filament assembly by their pyrin-domains (PYDs; e.g., viral nucleic acids, reactive oxygen species, specific lipids from damaged mitochondria, and disruption of the trans Golgi network) (Andreeva et al., 2021; Fernandes-Alnemri et al., 2009; Hornung et al., 2009; Iyer et al., 2013; Roberts et al., 2009; Zhong et al., 2013); PYDs are ∼100 amino-acid (a.a.) 6-helix bundle proteins that belong to the Death-Domain (DD) family often found in apoptotic and inflammatory signaling pathways (Park et al., 2007). The upstream PYD oligomers then induce the filament assembly by the PYD of the central adaptor ASC (ASC^PYD^), resulting in oligomerization/filamentation of its CARD (ASC: Apoptosis-associated-speck-forming-protein-containing-caspase-recruiting-domain (CARD)) (Broz & Dixit, 2016; Kagan et al., 2014; Lu et al., 2014; Lu & Wu, 2015). Finally, ASC^CARD^ oligomers recruit pro-caspase-1 and induce its filament assembly, activating the enzyme by proximity induced auto-proteolysis (Broz & Dixit, 2016; Kagan et al., 2014; Lu et al., 2014; Lu & Wu, 2015). Caspase-1 executes two key innate immune responses, namely the cleavage/maturation of pro-inflammatory cytokines such as interleukin-1β and -18, and the initiation of pyroptosis (Broz & Dixit, 2016; Kagan et al., 2014; Lu et al., 2014; Lu & Wu, 2015).

A hallmark of inflammasome assembly is its binary (on-or-off) nature (Cai et al., 2014; Franklin et al., 2014; Matyszewski et al., 2018; Shen et al., 2021). That is, once assembled, inflammasomes do not dissociate (Franklin et al., 2014; Matyszewski et al., 2018). Moreover, multiple positive feedback loops between upstream receptors and ASC not only bolster the assembly, but also result in prion-like self-perpetuation (Cai et al., 2014; Matyszewski et al., 2018). Such an inherently irreversible assembly mechanism in turn would necessitate extrinsic factors to prevent persistent activities. Indeed, mammalian pyrin-only-proteins (POPs) have emerged as major inhibitors of inflammasomes (de Almeida et al., 2015; Khare et al., 2014; Periasamy et al., 2017; Ratsimandresy et al., 2017), functioning analogous to CARD-only proteins (COPs) that interfere with the oligomerization/activation of pro-caspases (Devi et al., 2020; Indramohan et al., 2018; Lu et al., 2016). It has been proposed that the target specificities of POPs are dictated by their a.a. sequence homologies to inflammasome PYDs (Devi et al., 2020; Indramohan et al., 2018). For example, POP1 is most homologous to ASC^PYD^ (65% sequence identity; Figure 1- Figures Supplement 1A) and thought to directly inhibit the nucleation of the ASC^PYD^ filament (de Almeida et al., 2015). POP2 shares 68% sequence identity to the PYD of Nod-like-receptor containing a PYD-2 (NLRP2^PYD^; Figure 1- Figures Supplement 1B); the primary target of POP2 is thought to be ASC, but it is also implicated in inhibiting the Absent-in-melanoma-2 (AIM2) receptor (Periasamy et al., 2017; Ratsimandresy et al., 2017). Finally, POP3 is most similar to AIM2^PYD^ (67% sequence identity, Figure 1- Figures Supplement 1C) and targets AIM2-like receptors (ALRs, e.g., AIM2 and interferon inducible protein 16 (IFI16)) (Khare et al., 2014).

Inflammasome filaments are highly ordered supra-structures that entail at least two distinct steps for assembly: rate-limiting nucleation followed by elongation (Kagan et al., 2014; Lu et al., 2014; Lu & Wu, 2015; Matyszewski et al., 2018). Moreover, although the PYDs of upstream receptors do not display significant a.a. sequence homologies (Kagan et al., 2014; Lu et al., 2014; Lu & Wu, 2015), they all assemble into structurally congruent helical filaments and signal through the common ASC adaptor, suggesting a degenerate-code-like recognition mechanism (Hochheiser, Behrmann, et al., 2022; Kagan et al., 2014; Lu et al., 2014; Lu & Wu, 2015; Matyszewski et al., 2021; Shen et al., 2019). At present, little is known about how POPs selectively target and regulate the assembly of such diverse yet homologous supramolecular structures. This is because the current understanding on the mechanism of inhibition by POPs remains entirely inferred from indirect measurements and phenotypic outcomes (de Almeida et al., 2015; Khare et al., 2014; Periasamy et al., 2017; Ratsimandresy et al., 2017).

Here, by combining *in silico*, *in cellulo*, and *in vitro* methods, we delineate the target specificity and inhibition mechanisms of human POPs. We find that POP1 is a poor inhibitor of ASC but impedes the assembly of upstream receptor filaments instead (e.g., AIM2, IFI16, NLRP3, NLRP6). POP2 not only suppresses the nucleation of ASC, but also interferes with the assembly of multiple upstream receptors. Finally, in addition to potently suppressing the assembly of ALR and NLRP filaments (e.g., elongation of AIM2^PYD^), POP3 suppresses the nucleation of ASC. Our results indicate that a combination of favorable and strongly unfavorable interactions is necessary for POPs to inhibit the assembly of PYD filaments. Together, we propose that, instead of being dictated by a.a. sequence homology, degenerate-code-like target selection and inhibition mechanisms underpin the regulation of inflammasome assembly by POPs.

## Results

### Rosetta interface analyses suggest broad target specificities of POPs

Several inflammasome receptor PYDs signal through ASC^PYD^ although their primary a.a. sequences vastly differ (Broz & Dixit, 2016; Kagan et al., 2014; Lu & Wu, 2015). Such a functional redundancy amongst different PYDs in turn suggests that sequence homology may not dictate the target specificity of POPs. To elucidate how POPs recognize and regulate the assembly of different PYD filaments, we first implemented Rosetta-based *in silico* apporach that we had recently developed to define the directionality of the AIM2-ASC inflammasome (Matyszewski et al., 2021). Briefly, PYDs assemble into helical filaments in which each protomer provides six unique protein-protein interaction surfaces (e.g., Figure 1A and Figure 1 - Figure Supplement 2A-B, denoted as “Type” 1a/b, 2a/b, and 3a/b) (Lu et al., 2014; Lu & Wu, 2015). As we had done before (Matyszewski et al., 2021), we created a honeycomb-like sideview of PYD filaments in which the middle protomer makes all six required contacts for filament assembly (Figure 1 and Figure 1- Figure Supplement 2B). We then calculated Rosetta interface energies (ΔGs) for each PYD filament (e.g., Figure 1A, left), and also determined the ΔGs for POP•PYD interactions by replacing the center protomer with each POP (e.g., Figure 1A, three honeycombs on the right). Of note, we decided to conduct our *in silico* and subsequent biochemical experiments on tractable inflammasome components with well-known biological significances such as ASC, AIM2, IFI16, NLRP3, and NLRP6 (Broz & Dixit, 2016; Hochheiser, Behrmann, et al., 2022; Kerur et al., 2011; Lu et al., 2014; Lu & Wu, 2015; Matyszewski et al., 2018; Matyszewski et al., 2021; Morrone et al., 2015; Morrone et al., 2014; Shen et al., 2019).

**Figure 1.**
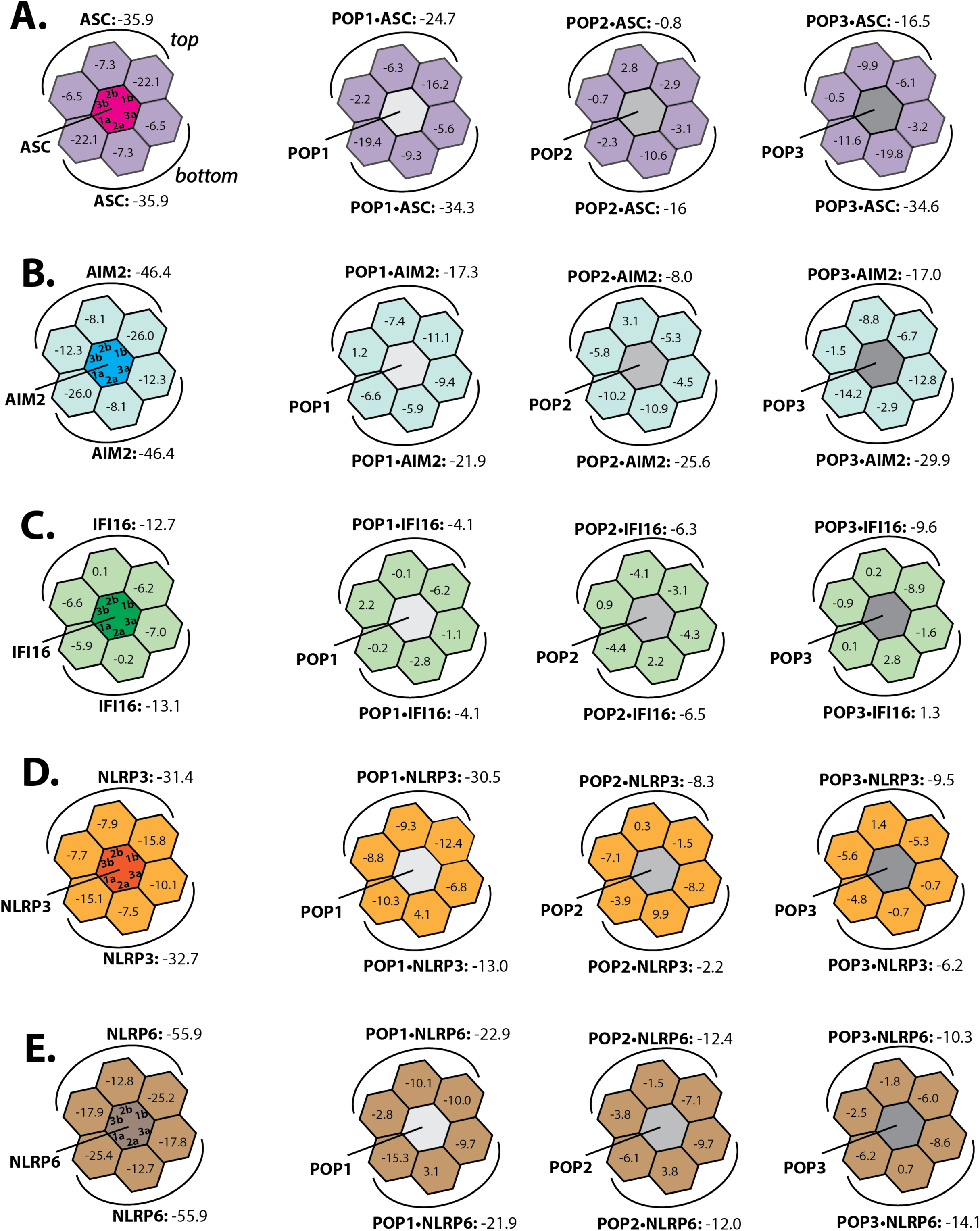
Rosetta *in silico* analyses of putative POP•PYD interactions. **(A-E)**. Rosetta interface energy scores (ΔGs) at individual filament interfaces for homotypic assemblies (left) and putative interactions with POPs (right). Each hexagon represents a PYD or POP monomer. The sum of ΔGs at the top and bottom half is also listed. The honeycombs were generated based on their respective cryo-EM structures except for IFI16^PYD^ whose filament structure is unknown. We generated a homology model of IFI16^PYD^ filament based on the GFP-tagged AIM2 filament (PDB: 6mb2), which produced more favorable ΔGs than the one generated from the tagless-AIM2^PYD^ filament (PDB: 7k3r; Figure 1 -Figure Supplement 2C).

We found previously that individual AIM2^PYD^ and ASC^PYD^ filaments assemble bi-directionally (i.e., extending from both top and bottom surfaces), with each pair of filament interface Types contributing symmetric ΔGs (e.g., both Type 1a and Type 1b surface show ΔG = -22.1 for ASC^PYD^•ASC^PYD^ in Figure 1A) (Matyszewski et al., 2021). The interface analysis results from other inflammasome PYDs also showed similar symmetric energy landscapes (Figure 1A-E, left), suggesting that bidirectional assembly is universal to all PYD filaments. We next noted that the overall ΔGs for individual PYD filaments were significantly more favorable than those from any POP•PYD interactions, which in turn suggested that excess POPs would be necessary to inhibit the assembly of inflammasome PYDs (e.g., the sum of ASC^PYD^•ASC^PYD^ interactions on the top half yields ΔG = -35.9, while that of POP1•ASC^PYD^ is -24.7; Figure 1A-E). POP1 showed more favorable overall ΔGs for ASC^PYD^ compared to POP2 or POP3 (Figure 1A), seemingly supporting the previous report (and sequency homology) that POP1 is the master regulator of ASC (de Almeida et al., 2015). However, although it was reported that POP2 can inhibit ASC^PYD^ more potently than POP1 *in vivo* (Ratsimandresy et al., 2017), its ΔGs were significantly less favorable (i.e., Figure 1A: POP1•ASC^PYD^=-24.7 vs. POP2•ASC = -0.8 on the top half). POP3, on the other hand, appeared to interact with ASC as favorably as POP1 at the bottom interfaces (Figure 1A; ΔG = -34.3 for POP1•ASC^PYD^ vs. ΔG =-34.6 for POP3•ASC^PYD^), suggesting that it could also inhibit the central adaptor. For AIM2^PYD^, all three POPs showed comparable overall ΔGs on the bottom half (Figure 1B), which suggested that each of them could target AIM2. IFI16 favored POP3 the most (Figure 1C; e.g., ΔG = -9.6 for POP3•IFI16^PYD^ on the top half); however, the other two POPs still showed more favorable ΔGs than homotypic IFI16^PYD^•IFI16^PYD^ interactions on at least one individual interface (Figure 1C; e.g., Type 2a for POP1, Type 2b for POP2 and Type 1b for POP3). These results in turn suggested that not only POP3, but POP1 and POP2 might also recognize IFI16. It has been speculated that POPs interfere with the recruitment of ASC by upstream receptors (de Almeida et al., 2015; Devi et al., 2020; Indramohan et al., 2018; Periasamy et al., 2017; Ratsimandresy et al., 2017); however, it remains unknown whether they do so by directly inhibiting NLRPs, which are the major class of inflammasome receptors. Our Rosetta analyses here suggest that POP1 could interact with NLRP3^PYD^ on the top half (Figure 1D; ΔG = -31.4 for NLRP3^PYD^•NLRP3^PYD^ vs. ΔG =-30.5 for POP1•NLRP3^PYD^), On the other hand, the interactions between POP2/3 and NLRP3^PYD^ appeared much less favorable (Figure 1D). Similar to NLRP3^PYD^, POP1 again showed more favorable energy scores toward NLRP6^PYD^ than POP2/3 (Figure 1E). Overall, although these results appear to rationalize some of the proposed target specificities of POPs (Devi et al., 2020; Indramohan et al., 2018), they also suggest a bit confounding interaction and recognition mechanisms, thus warranting further investigations via biochemical approaches.

### Mechanisms of ASC inhibition by POPs

Previous investigations on POP•PYD interactions have predominantly relied on *in cellulo* downstream signaling activities and *in vivo* phenotypes (de Almeida et al., 2015; Khare et al., 2014; Periasamy et al., 2017; Ratsimandresy et al., 2017). Although establishing the physiological relevance of POPs, these studies have left large voids in understanding their target selection and inhibition mechanisms. Because our *in silico* analyses did not immediately yield clear explanations for such questions, we set out to test POP•PYD interactions using more direct *in cellulo* and *in vitro* methods, focusing on ASC first. When ectopically expressed in HEK293T cells, C-terminally mCherry-tagged ASC^PYD^ forms filaments and full-length ASC (ASC^FL^) forms puncta (Matyszewski et al., 2021). Importantly, HEK293T cells do not contain any endogenous inflammasome components or POPs, providing an ideal cellular system for directly tracking their interactions (de Almeida et al., 2015; Matyszewski et al., 2021; Shi et al., 2016). Here, we first tested whether co-transfecting C-terminally eGFP-tagged POPs hinders the oligomerization of ASC. Surprisingly, compared to co-transfecting eGFP alone, POP1-eGFP minimally inhibited the filament assembly of ASC^PYD^-mCherry (≤ 20% suppression vs. eGFP control; Figure 2A-B). By contrast, co-transfecting POP2-eGFP or POP3-eGFP significantly reduced the amount of ASC^PYD^-mCherry filaments, with POP2 being more effective (Figure 2A-B; note more diffused mCherry signals and reduction in linear filaments in the presence of POP2 and POP3 in Figure 2A). Furthermore, POP1 reduced the number of ASC^FL^ puncta by ∼ 30%, yet POP2 and POP3 were again more effective in preventing punctum formation (up to ∼ 80% reduction, Figure 2C- D). These results suggest that POP1 may not directly target ASC^PYD^. Moreover, unlike COPs that co-assemble with CARDs into filaments (Lu et al., 2016), our results indicate that POPs suppress filament assembly altogether.

**Figure 2.**
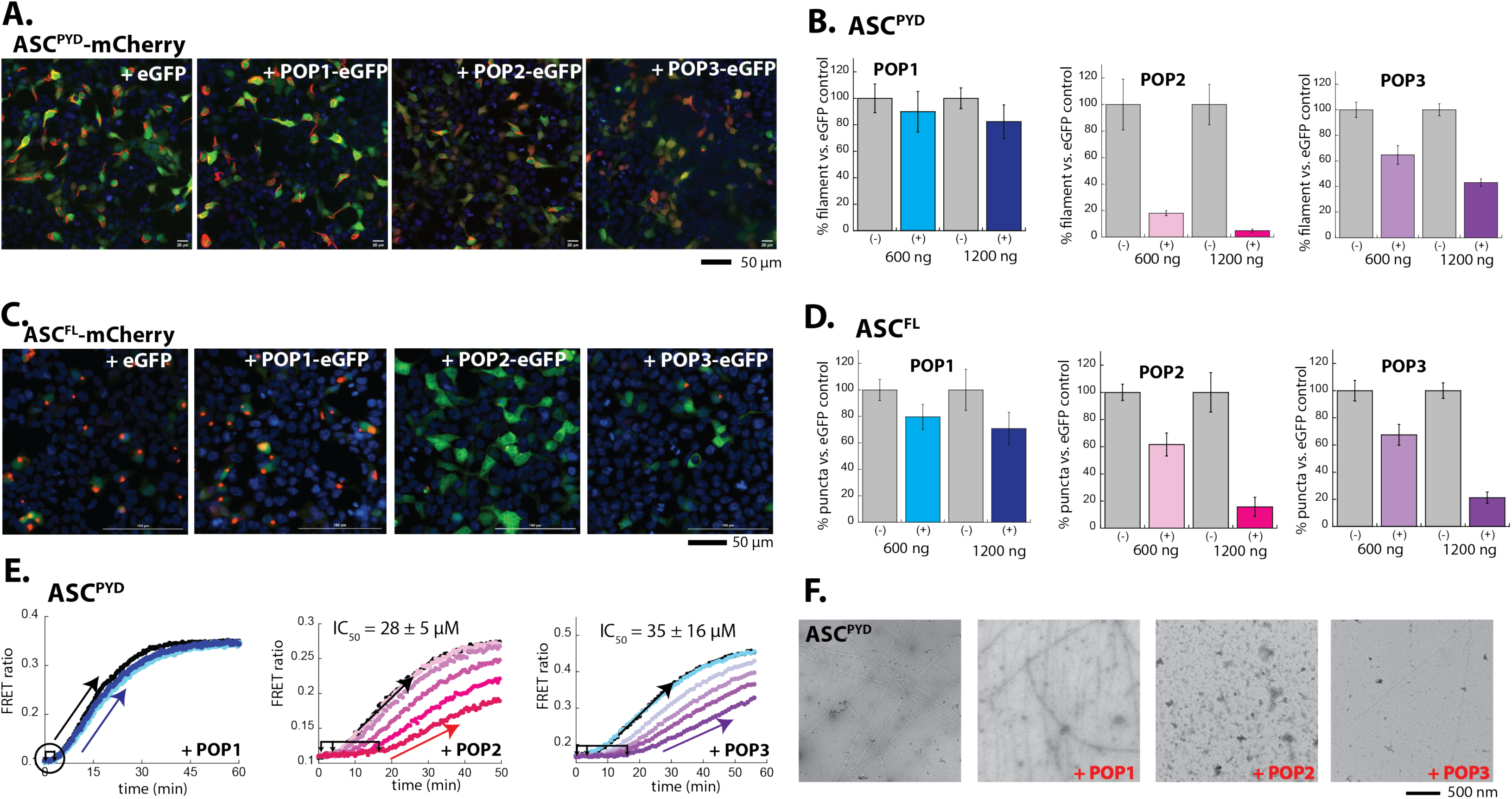
POP1 does not directly inhibit ASC. **(A)** Sample fluorescent microscope images of HEK293T cells co-transfected with mCherry- tagged ASC^PYD^ (300 ng; crimson) plus eGFP alone or eGFP-tagged POPs (1200 ng; green). Blue: DAPI. **(B)** The relative amounts of ASC^PYD^-mCherry filaments (300 ng plasmid) in HEK293T cells when co-transfected with POP-eGFP (+) or eGFP alone (-) (600 and 120 ng plasmids). *N* ≥ 4. **(C)** Sample fluorescent microscope images of HEK293T cells co-transfected with mCherry- tagged ASC^FL^ (300 ng; crimson) plus eGFP alone or eGFP-tagged POPs (1200 ng; green). Blue: DAPI. **(D)** The relative amounts of ASC^FL^-mCherry puncta (300 ng plasmid) in HEK293T cells when co-transfected with POP-eGFP (+) or eGFP (-) (600 and 120 ng plasmids). *n* ≥ 4. **(E)** Time-dependent increase in FRET signals of a donor- and acceptor-labeled ASC^PYD^ (2.5 µM total, black circle) was monitored with increasing concentrations POP1(50 and 150 µM), POP2 (3.3, 6.7, 13.3, 26.7, and 40 µM), or POP3 (3.25, 7.5, 15, 20, and 30 µM); darker shades correspond to increasing POP concentrations. Two- and three-headed arrows indicate the increase in apparent nucleation time (or lack thereof). Arrows pointing upper right directions indicate the change (or lack thereof) in the elongation phase in the presence of the highest POP concentrations used. Data shown are representatives of at least three independent measurements (IC_50_s are average values of these experiments. *n* = 3). **(F)** nsEM images of ASC^PYD^ filaments (2.5 µM) in the presence and absence of POP1 (150 µM), POP2 (40 µM), or POP3 (30 µM).

Our unexpected observations contradict the role of POP1 in directly inhibiting ASC assembly (de Almeida et al., 2015; Devi et al., 2020; Indramohan et al., 2018). To further test our *in cellulo* results, we then generated recombinant POPs to investigate their inhibitory mechanisms. POP1 behaved as a monomer without forming filaments or higher-order species in our hands (Figure 2- Figure Supplement 1A-B). On the other hand, recombinant POP2 and POP3 were prone to aggregation/degradation during purification and required an N-terminal maltose binding protein (MBP) tag to obtain intact proteins (Figure 2- Figure Supplement 1A). Cleaving MBP via Tobacco Etch Virus protease (TEVp) indicated that POP2 and POP3 form elongated oligomers with undefined structures (Figure 2- Figure Supplement 1B).

We incorporated recombinant POPs into our well-established polymerization assay in which we track the Förster Resonance Energy Transfer (FRET) ratio between a 1:1 mixture of donor- and acceptor labeled PYDs (Matyszewski et al., 2018; Matyszewski et al., 2021; Mazanek & Sohn, 2019); the auto-assembly of individual PYDs is suppressed by an N-terminal MBP tag and polymerization is triggered by cleaving MBP via TEVp. Of note, our assay displays two distinct phases of filament assembly, namely the rate limiting nucleation (initial lag; double/triple-headed arrows in Figure 2E) followed by elongation (exponential/linear phase; single-headed arrows pointing to the upper righthand corner in Figure 2E (Matyszewski et al., 2018; Matyszewski et al., 2021; Mazanek & Sohn, 2019).

Consistent with our cellular imaging assays (Figure 2A-B), POP1 did not affect the oligomerization kinetics of ASC^PYD^ up to the highest concentration we could achieve (Figure 2E; 150 µM POP1 vs. 2.5 µM ASC^PYD^). Moreover, the presence of POP1 did not affect the formation of ASC^PYD^ filaments when visualized with nsEM (Figure 2F). On the other hand, both POP2 and POP3 significantly prolonged the initial lag phase of ASC^PYD^ polymerization in a dose dependent manner (up to ∼ 15 min delay in nucleation; Figure 2E), while also moderately affecting the elongation phase (≤ 20% reduction in the slope Figure 2E). Estimating the polymerization half-times at different POP concentrations indicated that POP2 and POP3 are similarly effective in suppressing the oligomerization of ASC^PYD^ (Figure 2E, IC_50_s). When visualized via nsEM, no ASC^PYD^ filaments were detected in the presence of POP2, and only a few filaments were seen with POP3 (Figure 2F). These results suggest that even if ASC^PYD^s form oligomers (rise in FRET signals in Figure 2E), most of them fail to assemble into intact filaments in the presence of POP2/3. Moreover, the complete absence of any ASC^PYD^ filaments in the presence of POP2 is consistent with our cellular experiments in which POP2 was most potent in inhibiting the central adaptor (Figures 2B and 2F). Together, our *in cellulo* and *in vitro* experiments consistently indicate that POP1 is only marginally effective in directly suppressing the polymerization of ASC. We also find that both POP2 and POP3 impede the nucleation of the ASC^PYD^ filament.

### Re-examining the Rosetta analyses in light of biochemical experiments

Our biochemical experiments indicated that excess POPs are required to inhibit the polymerization of ASC (Figure 2E), which is in agreement with the Rosetta analyses in which no POP•PYD interactions were more favorable than homotypic PYD•PYD interactions (Figures 1 and 2). However, although our *in silico* analyses suggested that POP1 should interact most favorably with ASC^PYD^ (Figure 1A), our *in vitro* and *in cellulo* experiments consistently showed that POP1 is only marginally inhibitory (Figure 2). We thus re-examined our Rosetta results in light of our biochemical experimental results, and noted that the interface energy profiles of POP2•ASC^PYD^ and POP3•ASC^PYD^ are different from that of POP1•ASC^PYD^. For instance, all three POPs contain favorable protein•protein interaction surfaces for ASC^PYD^ (ΔΔG = ΔG^PYD•PYD^- ΔG^POP•PYD^ ≤ 3.5; arbitrarily determined, marked as blue dots in Figure 2- Figure Supplement 2). However, although both POP2 and POP3 show multiple interfaces with significantly unfavorable ΔGs than ASC^PYD^•ASC^PYD^ interactions, POP1•ASC^PYD^ interfaces lack such negative interactions (ΔΔG ≥ 10.0; marked as red dots in Figure 2-Figure Supplement 2). These observations in turn raised the hypothesis that a combination of favorable (recognition) and unfavorable interfaces (repulsion) is necessary for POPs to interfere with the assembly of inflammasome PYDs.

### Mechanisms of ALR inhibition by POPs

We then set out to test our amended interpretation of Rosetta analyses on other POP•PYD interactions such as those with AIM2 and IFI16. Here, we noted a mixture of both favorable and unfavorable interactions between all three POPs and both ALRs (Figure 3-Figure Supplement 1A-B), raising the possibility that not only POP3, but the other two POPs might also inhibit the oligomerization of ALRs. To test this, we monitored the filament assembly of AIM2^PYD^- mCherry and IFI16^PYD^-mCherry in HEK293T cells (Figure 3A-D). Compared to co-transfecting with eGFP alone, POP1-eGFP reduced the number of AIM2^PYD^ and IFI16^PYD^ filaments, apparently more effectively than against ASC^PYD^ (Figure 2B vs. Figures 3B and 3D; e.g., at 1200 ng POP1, AIM2^PYD^ and IFI16^PYD^ assemblies were inhibited ∼60%, while ASC^PYD^ assembly was suppressed ∼20%). On the other hand, co-transfecting POP2 or POP3 essentially obliterated the filament assembly of AIM2^PYD^ and IFI16^PYD^ (Figures 3A-B and 3C-D). In our FRET assays tracking AIM2^PYD^ polymerization, all three POPs decreased the slope of the linear phase in a dose dependent manner without significantly affecting the initial lag phase (Figure 3E); recombinant IFI16^PYD^ does not form filaments in our hands (Morrone et al., 2014). Our observations indicate that all three POPs can interfere with the elongation of the AIM2^PYD^ filament, with POP3 being most effective (Figure 3E, IC_50_s). Moreover, imaging AIM2^PYD^ filaments using nsEM in the presence of POP1 revealed that the filaments are shorter and fewer, and the presence of POP2/3 abrogated filament formation (Figure 3F). As seen from ASC^PYD^, the dearth of filaments in the presence of POP2/3 in our nsEM and *in cellulo* experiments (Figure 3B, D and F) indicated that AIM2^PYD^ oligomers rarely progressed into functional filaments (i.e., rise in FRET signals in Figure 3E vs. the lack of filaments in Figures 3F).

**Figure 3.**
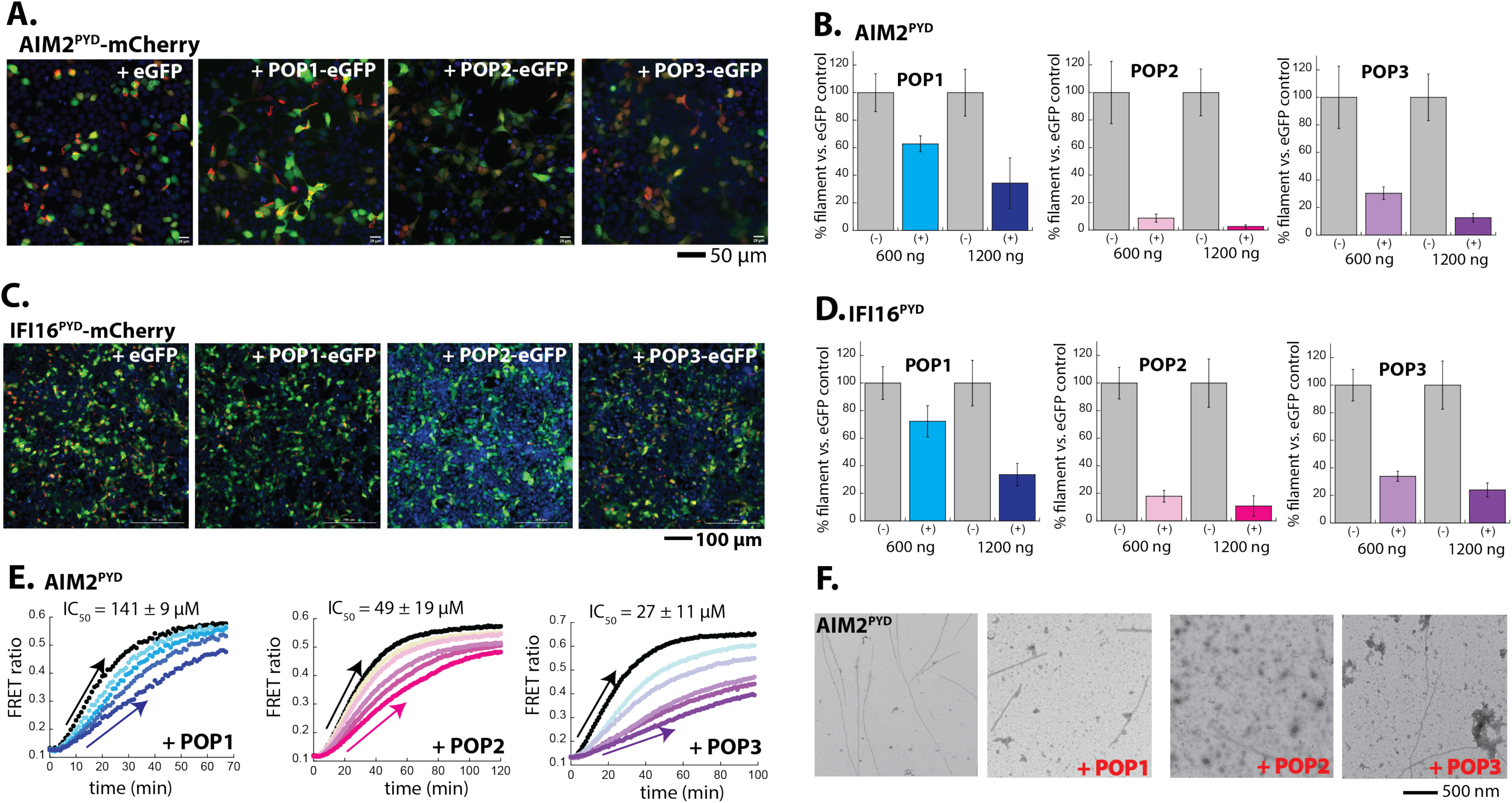
Inhibition of ALR assembly by POPs. **(A)** Sample fluorescent microscope images of HEK293T cells co-transfected with mCherry- tagged AIM2^PYD^ (300 ng; crimson) plus eGFP alone or POP-eGFP (1200 ng; green). Blue: DAPI. **(B)** The relative amounts of AIM2^PYD^-mCherry filaments (300 ng plasmid) in HEK293T cells when co-transfected with POP-eGFP (+) or eGFP alone (-) (600 and 120 ng plasmids). *n* ≥ 4. **(C)** Sample fluorescent microscope images of HEK293T cells co-transfected with mCherry- tagged IFI16^PYD^ (300 ng; crimson) plus eGFP alone or eGFP-tagged POPs (1200 ng; green). Blue: DAPI **(D)** The relative amount of IFI16^PYD^-mCherry filaments (300 ng plasmid) in HEK293T cells when co-transfected with POP-eGFP (+) or eGFP alone (-) (600 and 120 ng plasmids). *N* ≥ 4. **(E)** Time-dependent increase in FRET signals of a donor- and acceptor-labeled AIM2^PYD^ (2.5 µM, black) was monitored with increasing concentrations of POP1 (25, 50, 100 and 150 µM), POP2 (12.5, 25, 50, and 75 µM), or POP3 (7.5, 15, 30, 40, and 50 µM); darker shades correspond to increasing POP concentrations. Arrows pointing upper right directions indicate the change (or lack thereof) in the elongation phase in the presence of the highest POP concentrations used. Data shown are representatives of at least three independent measurements (IC_50_s are average values of these experiments. *n* = 3). **(F)** nsEM images of AIM2^PYD^ filaments (2.5 µM) in the presence and absence of POP1 (100 µM), POP2 (50 µM), or POP3 (40 µM).

It is noteworthy that the oligomerization of PYD is important for stable dsDNA binding by ALRs (Morrone et al., 2015; Morrone et al., 2014). Conversely, although isolated AIM2^PYD^ can form filaments by mass-action (i.e., high concentrations; (Morrone et al., 2015)), dsDNA provides a one-dimensional diffusion scaffold to facilitate the assembly of full-length ALRs at significantly lower concentrations (Morrone et al., 2015; Morrone et al., 2014). AIM2^FL^ forms punctum-like oligomers when transfected in HEK293T cells (Matyszewski et al., 2021), and we found that POP1 slightly reduced the number of AIM2^FL^ puncta, while POP2 and POP3 were more effective (Figure 3- Figure Supplement 1C-D); IFI16^FL^ localizes in the nucleus (Antiochos et al., 2018; Kerur et al., 2011; Li et al., 2012), precluding our investigation with cytosolic POPs (Figure 3-Figure Supplement 1E). Consistent with the lack of significant inhibition in cells (Figure 3- Figure Supplement 1 C-D), POP1 failed to interfere with dsDNA-binding/oligomerization of recombinant AIM2^FL^ (Figure 3- Figure Supplement 2A). However, POP2 and POP3 still inhibited the dsDNA binding of AIM2, while only POP3 was inhibitory toward the dsDNA binding of recombinant IFI16^FL^ (Figure 3- Figure Supplement 2B). Together, our observations indicate that POP3 directly inhibits ALR assembly (elongation in particular for AIM2^PYD^). We also find that POP1 and POP2 can inhibit the assembly of ALR filaments, with POP2 being significantly more effective than the former; the presence of activating ligands can diminish the inhibitory effect of POPs (Figure 3- Figure 3 Supplement 2). Furthermore, these results are consistent with our amended interpretation of Rosetta analyses in which a combination of favorable and unfavorable interfaces allow POPs to target and inhibit PYD filament assembly.

### POP1 likely targets upstream receptors instead of ASC

Although it has been speculated that POP1 and POP2 would interfere with the recruitment of ASC by the NLRP receptors (de Almeida et al., 2015; Devi et al., 2020; Indramohan et al., 2018; Periasamy et al., 2017; Ratsimandresy et al., 2017), it remains unknown whether either POP can directly suppress the filament assembly of NLRP^PYD^s. Of note, our investigations here revealed that POP1 is ineffective in inhibiting the oligomerization of ASC (Figure 2). Moreover, albeit less inhibitory than POP2 or POP3, POP1 was more effective in suppressing the assembly of AIM2^PYD^ and IFI16^PYD^ filaments than that of ASC^PYD^ (Figure 3). These observations prompted the new hypothesis that the role of POP1 is to interfere with the assembly of upstream receptors rather than directly inhibiting ASC (i.e., a “decoy” ASC; targeting ASC or multiple upstream receptors would result in the same phenotype). Indeed, our Rosetta analyses indicated that POP1 can make a combination of favorable and unfavorable interactions with both NLRP3^PYD^ and NLRP6^PYD^ (Figure 4- Figure Supplement 1). Furthermore, POP2 and POP3 also showed favorable and unfavorable interactions with NLRP3 (Figure 4- Figure Supplement 1A); although the ΔGs between NLRP6 and POP2/3 were largely unfavorable, the Type 3a surface showed an energy score that might allow the two POPs to recognize NLRP6 if present at high enough concentrations (ΔG∼ -9; Figure 4- Figure Supplement 1B, marked as light pink); our reasoning is based on ΔGs ∼ -9 seen from native PYD•PYD interactions (e.g., NLRP3^PYD^•NLRP3^PYD^ shows ΔGs of ∼ -8 and -10 on Type 2 and Type 3 interfaces, Figure 1 and Figure 4- Figure Supplement 1A).

The activation mechanisms of NLRPs are complex and involve different types of active and inactive oligomers (Andreeva et al., 2021; Gong et al., 2021; Hochheiser, Pilsl, et al., 2022; Lu et al., 2014; Ohto et al., 2022; Sharif et al., 2019; Sharma & de Alba, 2021; Shen et al., 2021; Shen et al., 2019), we thus decided to monitor whether POPs impede the filament assembly using the isolated PYDs of NLRP3 and NLRP6, which formed filaments when ectopically expressed in HEK293T cells (Figure 4A/C). Of note, NLRP2^PYD^-mCherry did not form filaments when expressed in HEK293T cells (Figure 4- Figure Supplement 1C), precluding further investigations despite its high sequence similarity to POP2 (Indramohan et al., 2018; Periasamy et al., 2017; Ratsimandresy et al., 2017). When co-transfected, POP1 was more effective in suppressing the filament assembly by both NLRP3^PYD^ and NLRP6^PYD^ than that of ASC^PYD^ (Figure 4 A-D). For example, with 1200 ng POP1, ASC^PYD^ assembly was only suppressed by ∼20%, but the filament assembly by NLRP3^PYD^ and NLRP6^PYD^ was suppressed 70% and 60%, respectively (Figure 2B vs. Figure 4B/D). On the other hand, POP2 was less effective in inhibiting the polymerization of NLRP^PYD^s than that of ASC^PYD^ (e.g., at 600 ng POP2, ASC^PYD^ assembly was abolished, but the assembly of NLRP3^PYD^ and NLRP6^PYD^ was barely suppressed; Figure 2B vs. Figure 4B/D). POP3 was almost equally effective in suppressing the polymerization of NLRP^PYD^s and ASC^PYD^, but not nearly as effectively as against ALRs (Figure 4A-D). For example, at 600 ng POP3, AIM2^PYD^ assembly was suppressed 70%, but those of ASC^PYD^ and NLRP^PYD^s were reduced by 40-50%; Figure 3B vs. Figures 2B, 4B, and 4D).

**Figure 4.**
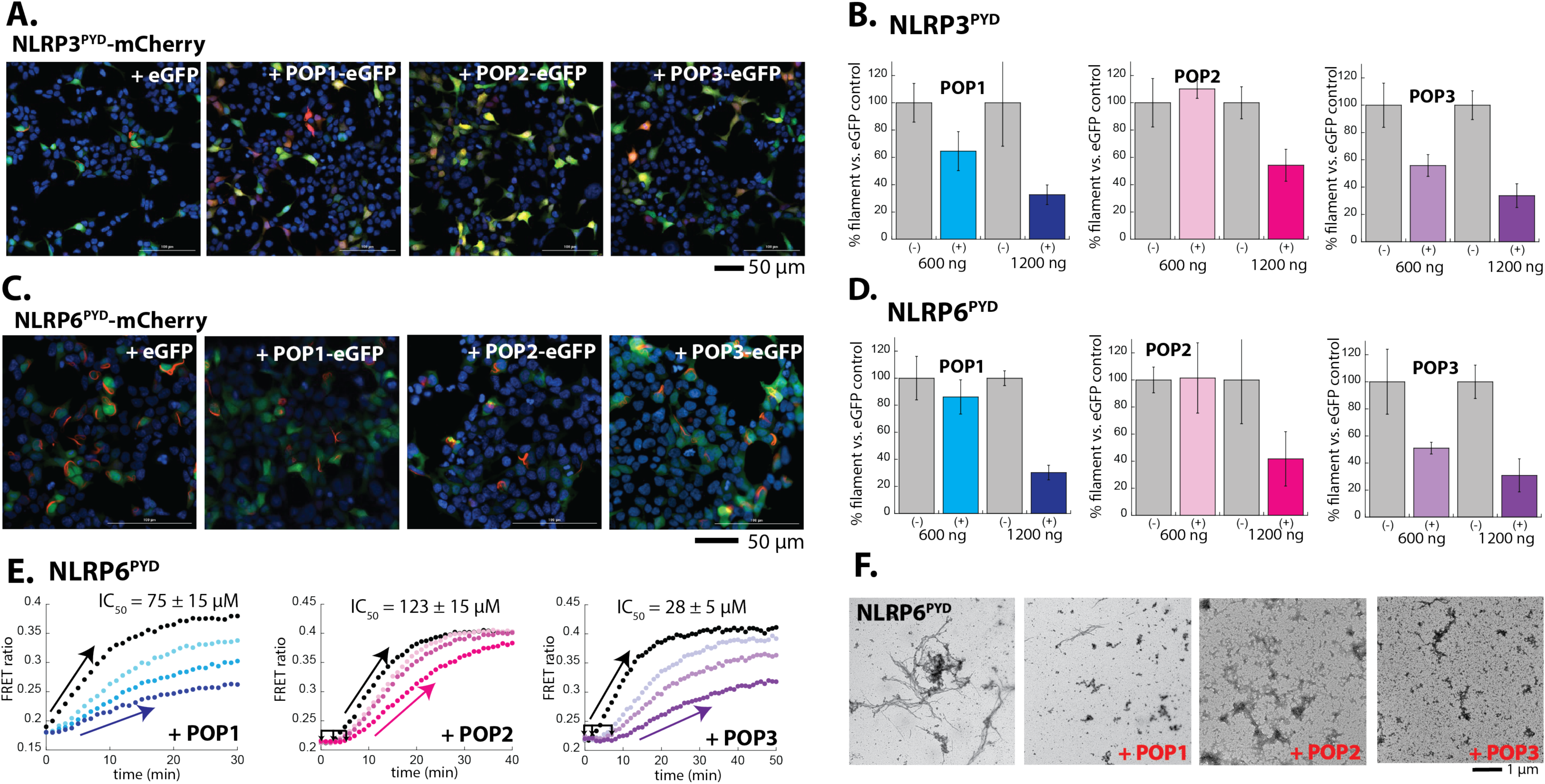
Inhibition of NLRP^PYD^ assembly by POPs. **(A)** Sample fluorescent microscope images of HEK293T cells co-transfected with mCherry- tagged NLRP3^PYD^ (600 ng; crimson) plus eGFP alone or eGFP-tagged POPs (1200 ng; green). Blue: DAPI **(B)** The relative amounts of NLRP3^PYD^-mCherry filaments (600 ng plasmid) in HEK293T cells when co-transfected with POP-eGFP (+) or eGFP alone (-) (600 and 120 ng plasmids). *n* ≥ 4. **(C)** Sample fluorescent microscope images of HEK293T cells co-transfected with mCherry- tagged NLRP6^PYD^ (300 ng; crimson) plus eGFP alone or eGFP-tagged POPs (1200 ng; green). Blue: DAPI **(D)** The relative amounts of NLRP6^PYD^-mCherry filaments (300 ng plasmid) in HEK293T cells when co-transfected with POP-eGFP (+) or eGFP alone (-) (600 and 120 ng plasmids). *n* ≥ 4. **(E)** Time-dependent increase in FRET signals of a donor- and acceptor-labeled NLRP6^PYD^ (2.5 µM, black) was monitored with increasing concentrations of POP1 (25, 50, and 100 µM). POP2 (30, 60, and 120 µM). POP3 (15, 30, and 60 µM); darker color shades correspond to increasing POP concentrations. Two and three-headed arrows indicate the increase in apparent nucleation time. Arrows pointing upper right directions indicate the change (or lack thereof) in the elongation phase in the presence of the highest POP concentrations used. Data shown are representatives of at least three independent measurements (IC_50_s are average values of these experiments. *n* = 3). **(F)** nsEM images of NLRP6^PYD^ filaments (5 µM) in the presence and absence of POP1 (100 µM), POP2 (30 µM), or POP3 (30 µM).

Next, using FRET-donor and acceptor labeled NLRP6^PYD^s, we then monitored whether POPs suppress the nucleation and/or elongation (recombinant NLRP3^PYD^ does not auto-assemble into filaments in our hands (e.g., (Bae & Park, 2011)). POP1 predominantly inhibited the elongation of NLRP6^PYD^ filament, and POP2 appeared to interfere with its nucleation. Although POP3 mostly reduced the elongation kinetics of NLRP6^PYD^, it also seemed to interfere with nucleation (Figure 4E). Consistent with these observations, the number and length of NLRP6^PYD^ filaments were significantly reduced in the presence of POPs (Figure 4F). The lack of filaments (Figure 4F) despite the increase in FRET signals (Figure 4E) again indicate that NLRP6^PYD^ oligomers fail to form intact filaments. Together with the results form investigating POP•ALR interactions, our observations are consistent with the idea that POP1 acts as a decoy ASC, interfering with the assembly of upstream PYDs. Moreover, our results further support the idea that favorable and unfavorable interactions are necessary for POPs to inhibit the assembly of inflammasome PYDs.

## Discussion

### Redefining the target specificity and inhibitory mechanisms of POPs

A hallmark of inflammasomes is their exceptional stability. For example, ASC is known to promote “solidification” of inflammasomes in a prion-like manner, allowing them to perpetuate even after cells undergo pyroptosis (Cai et al., 2014; Franklin et al., 2014; Shen et al., 2021). Moreover, AIM2 and IFI16 filaments also persist and are even stigmatized as autoantigens in debilitating autoimmune disorders such as systemic lupus erythematosus (SLE) and Sjögren’s Syndrome (Antiochos et al., 2018; Antiochos et al., 2022; Baer et al., 2016). Indeed, persisting inflammasome oligomers and their aberrant activities are implicated in a wide range of human diseases even including COVID-19 (Karki et al., 2017; Tartey & Kanneganti, 2020; Vora et al., 2021). Thus, it is critical for the host to carefully modulate the assembly of inflammasomes at the onset, as it would be much more difficult to demolish such hyper-stable supra-structures. POPs have emerged as major endogenous regulators of inflammasomes by directly interfering with the assembly of PYD filaments, functioning analogous to COPs that target the oligomerization of pro-caspases (Devi et al., 2020; Indramohan et al., 2018). Although the biological significances of POPs are well-established (de Almeida et al., 2015; Devi et al., 2020; Indramohan et al., 2018; Khare et al., 2014; Periasamy et al., 2017; Ratsimandresy et al., 2017), their intrinsic target specificities and inhibition mechanisms have remained speculative. To address these issues, we employed tools and methods that allow us to directly probe the protein•protein interactions between human PYDs and POPs.

Our investigations here reveal that POPs interfere with the polymerization (nucleation and/or elongation) of various inflammasome filaments without co-assembling, which is different from COPs that co-assembles into filaments with CARD of caspase-1 (Lu et al., 2016). Moreover, also unlike COPs that can inhibit the assembly of pro-caspases at sub-stoichiometric concentrations (Lu et al., 2016), excess POPs were necessary to inhibit inflammasome PYDs, especially when an activating ligand was present (dsDNA for AIM2 and IFI16; Figure 3- Figure supplement 2). Additionally, POP2/3 largely suppressed the nucleation of ASC (e.g., prolonged lags in Figure 2E), yet these POPs also interfered with the elongation of upstream receptors (e.g., Figures 3E and 4E). Although often considered as a harmful phenomenon, inflammation is integral to host innate defense and survival (Bennett et al., 2018; Meizlish et al., 2021). We reason that being able to modulate two key assembly steps (nucleation and elongation) while requiring excess POPs is well suited for effectively regulating inflammasome activities without impairing their proper functions.

Although it was previously proposed that POP1 would directly interact with ASC^PYD^, such conclusions were indirectly deduced from using *in cellulo* and *in vivo* systems that harbor multiple inflammasome receptors and regulators (de Almeida et al., 2015). Here, we provide an important addendum to the inhibitory mechanism of POP1. That is, POP1 is minimally effective in inhibiting the polymerization of ASC^PYD^. However, it is likely to modulate inflammasomes through interfering with the oligomerization of upstream PYDs, most notably by halting their elongation (e.g., Figure 3E and 4E). Still, our results indicate that POP1 is less potent than the other two POPs in inhibiting inflammasomes (Figures 2-4). It is noteworthy that the expression of POP1 is predominantly induced by IL-1β (de Almeida et al., 2015), a major final product of inflammasome cascades (Broz & Dixit, 2016; Zheng et al., 2020). Thus, it is likely that POP1 is part of a negative feedback loop for attenuating the signaling activity of inflammasomes. Also of note, the role of negative feedback by definition is to provide stability to a given system, but not to terminate it (Brandman & Meyer, 2008). Thus, in our view, POP1 is ideally positioned to generate well-balanced inflammasome responses by suppressing the excessive elongation (perpetuation) of upstream receptor filaments (Figure 5).

**Figure 5.**
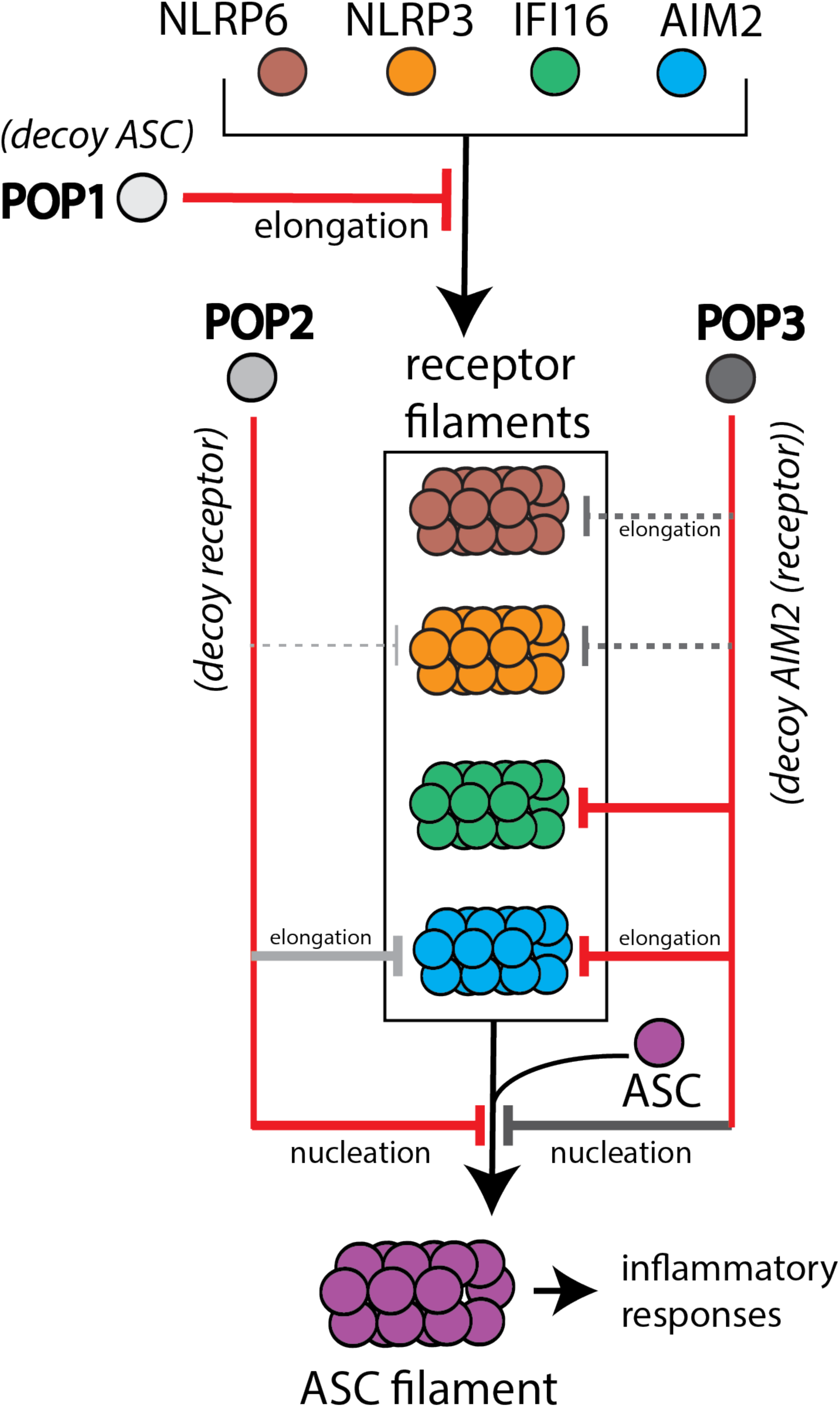
Target selection and mode of inhibition of POPs. A cartoon summarizing the refined intrinsic target specificity of POPs. Solid red lines indicate the most likely primary inhibitory targets for each POP. Solid gray lines indicate an additional target that each POP could also directly inhibit. Dotted gray lines indicate other possible targets for each POP.

POP2 is thought to have a similar function as POP1 by suppressing the activation of ASC (Periasamy et al., 2017; Ratsimandresy et al., 2017). Interestingly, despite the lack of sequence similarity to ASC^PYD^, POP2 is reportedly more effective than POP1 in preventing spurious inflammasome activities *in vivo* (Periasamy et al., 2017; Ratsimandresy et al., 2017). Nevertheless, it has remained unclear whether it directly suppresses ASC assembly or those of upstream receptors. We find here that POP2 can potently suppress the nucleation of ASC^PYD^ (Figure 2). Additionally, although less effective toward NLRP^PYD^s (Figure 4), POP2 also suppressed the assembly of AIM2^PYD^ and IFI16^PYD^, indicating that it has intrinsic capability to inhibit upstream receptors (Figures 3-4). Of note, while POP2 interfered with dsDNA-mediated assembly/binding of AIM2^FL^, it failed to inhibit that of IFI16^FL^ (Figure 3- Figure Supplement 1B and 2). These observations collectively support the idea that POP2 directly targets ASC assembly while simultaneously inhibiting upstream receptors such as AIM2 (Periasamy et al., 2017). We reason that the presence of dsDNA would allow IFI16 (and AIM2 to some extent) to outcompete any POP•PYD interactions to properly initiate their respective inflammatory pathways (Figure 3- Figure Supplements 1C/D and 2), again suggesting that the role of POPs is to regulate inflammasomes, but not to shut them down altogether. It is noteworthy that the expression of POP2 is readily induced by a wide variety of pro- and anti-inflammatory cytokines (Ratsimandresy et al., 2017). Considering that it primarily targets ASC, we postulate that POP2 would act as the pan-inflammasome inhibitor essential for preventing spurious innate immune responses (Figure 5).

POP3 was identified as a selective inflammasome inhibitor targeting ALRs (Khare et al., 2014). In addition to inhibiting the elongation of the AIM2^PYD^ filament, we find here that POP3 can robustly inhibit the nucleation of ASC^PYD^ (Figure 2), revealing its dual targeting function similar to POP2. We also find that, in principle, POP3 can inhibit the assembly of NLRP^PYD^s (Figure 4). Unlike the other two POPs, POP3 is exclusively induced by interferons (IFNs) (Khare et al., 2014), a major cytokine family that counteracts IL-1β (Mayer-Barber & Yan, 2017). Considering that IFNs also drive the expression of ALRs (Antiochos et al., 2018; Khare et al., 2014), we postulate that although POP3 is intrinsically capable of inhibiting various inflammasome receptors, the contextual expression would dictate its *in vivo* targets (Figure 5).

Although the PYDs of upstream inflammasome receptor do not share sequence homologies with ASC^PYD^ or even with one another, they all signal through ASC^PYD^ (Broz & Dixit, 2016; Lu & Wu, 2015). Of note, such a hub-like function of ASC suggests a degenerate genetic-code-like recognition mechanism (i.e., various a.a. sequences (and combinations) at the recognition interfaces are tolerated and accepted between ASC^PYD^ and receptor PYDs) (Matyszewski et al., 2021). Analogously, our observations here indicate that sequence homology does not dictate the intrinsic target specificity of POPs. Instead, we find that each POP has intrinsic affinity toward multiple inflammasome components. Moreover, our results suggest that POPs act as a decoy receptor and/or adaptor (Figure 5). For example, POP1, which is most similar to ASC (Figure 1 - Figure Supplement 1A), acts as a decoy adaptor for upstream receptors (Figure 5). POP2, which is most similar to NLRP2^PYD^ (Figure 1 -Figure Supplement 1B), predominantly acts as a decoy receptor for ASC, suppressing its nucleation (Figure 5). Finally, our results show that POP3, which is homologous to AIM2^PYD^ (Figure 1 -Figure Supplement 1C), not only inhibits the assembly of ALRs, but also can act as a decoy receptor to suppress the nucleation of ASC^PYD^ (Figure 5). Although laboratory practices can selectively activate different inflammasome receptors, pathogen invasion and various inflammatory conditions provide activation signals for multiple inflammasome receptors (Broz & Dixit, 2016; Zheng et al., 2020). Thus, we propose that multifaceted target specificities and inhibition mechanisms (nucleation and/or elongation) by POPs would provide greater assurances for preventing spurious inflammasome activities.

### Design principles for inflammasome inhibition by POPs

Our use of *in silico* simulations has provided a solid foundation for subsequent biochemical investigations. For instance, consistent with our Rosetta analyses showing that homotypic PYD•PYD interactions are preferred in all cases, excess POPs were required to suppress the assembly of inflammasome PYDs (e.g., Figures 2E, 3E, and 4E). Moreover, although somewhat counterintuitive at first, further examining Rosetta *in silico* results in light of our initial biochemical experiments suggested that a combination of favorable and unfavorable interactions underpin the target specificity of POPs. For example, POP1•ASC^PYD^ lacked any strong unfavorable interactions (Figure 2- Figure Supplement 2), resulting in failure to inhibit the polymerization of ASC^PYD^ (Figure 2). On the other hand, although the POP1•NLRP3^PYD^ interfaces displayed only one strong unfavorable interface coupled with multiple favorable interactions (Figure 4- Figure Supplement 1A), POP1 was still able to inhibit the assembly of NLRP3^PYD^ filaments (Figure 4A-B). Surprisingly, NLRP6^PYD^ was inhibited by POP2 and POP3 despite the lack of any clear favorable interactions (Figure 4E-F and Figure 4 Figure Supplement 1B), which is also in contrast to POP1•ASC^PYD^ interactions where the absence of any strong unfavorable interactions was correlated with the lack of inhibition (Figure 2 and Figure 2 Figure Supplement 2). PYD filaments are highly organized helical structures- all six interfaces from each protomer are necessary to assemble into intact, functional PYD filaments (Hochheiser, Behrmann, et al., 2022; Lu et al., 2014; Matyszewski et al., 2021; Shen et al., 2019). We postulate that even weakly favorable interactions at one surface (e.g., Type 3a between POP2/3 and NLRP6^PYD^) would be sufficient for POPs associate with their targets and interfere with filament assembly via any unfavorable interactions at the other five available surfaces.

We envision that the nonequilibrium assembly of inflammasome filaments plays a major role in defining the mechanism of regulation by POPs (i.e., kinetically driven without having conceivable off-rates) (Cai et al., 2014; Matyszewski et al., 2018). For instance, considering that all POP and PYD protomers share the same overall structure (shape complementarity), we reason that any (semi) favorable protein-protein interaction interfaces would allow POPs and PYDs to associate transiently (i.e., classic reversible protein•protein interaction equilibrium). However, the more favorable homotypic PYD•PYD interactions would readily outcompete such meta- stable interactions especially when only basal amounts of POPs are present (Figure 5- Figure Supplement 1A). Moreover, as indicated from the failure to inhibit ASC^PYD^ by POP1, if POPs do not contain any strong unfavorable interactions against target PYDs, homotypic PYD•PYD interactions would still outcompete even excess POPs and lock themselves into irreversible filament assembly (Figure 5- Figure Supplement 1B). However, when such meta-stable POP•PYD complexes contain at least one very unfavorable interface, excess POPs would then expose a multitude of adverse protein•protein interaction surfaces that would hamper filament assembly (Figure 5- Figure Supplement 1C). Also of note, given that POP2 and POP3 can form oligomers (Figure 2- Figure Supplement 1B-C), it is highly likely that multimeric POPs are more effective in preventing the association of inflammasome PYDs (Figure 5- Figure Supplement 1C). We found previously that the recognition between AIM2 and ASC occurs when at least one is filamentous (Matyszewski et al., 2021). Thus, it is also possible that POPs might preferentially interact with oligomeric PYDs that are not yet fully filamentous (e.g., (pseudo)-nucleation unit), trapping them into non-functional states (Figure 5- Figure Supplement 1C). Future investigations using molecular dynamics simulations and extensive mutagenesis will further delineate the complexity of oligomerization mechanisms and target specificities of POPs in more detail. Overall, our multi-disciplinary approach provides an example of how to use *in silico* predictions judiciously for investigating multipartite protein-protein interactions.

## Materials and Methods

### Rosetta Simulation

The InterfaceAnalyzer script in Rosetta was used to determine the interaction energy at individual interfaces of the honeycomb (Matyszewski et al., 2021). We used the cryo-EM structures of ASC^PYD^ (PDB: 3j63; (Lu et al., 2014)), AIM2^PYD^ (PDB: 7k3r; (Matyszewski et al., 2021)), NLRP3^PYD^ (PDB: 7pdz; (Hochheiser, Behrmann, et al., 2022)) and NLRP6^PYD^ (PDB: 6ncv; (Shen et al., 2019)) filaments to generate corresponding honeycombs. Because the structure of the IFI16^PYD^ filament is unknown, we used the eGFP-AIM2^PYD^ filament (PDB: 6mb2; (Lu et al., 2015)) that shows a pentameric filament base as a template; using the untagged AIM2^PYD^ filament (PDB: 7k3r; (Matyszewski et al., 2021)), which shows a hexameric filament base, as a template resulted in largely unfavorable energy scores (Figure 1- Figure Supplement 1C). For POPs, we used the crystal structure of POP1 (PDB: 4qob), and generated homology models of POP2 and POP3 based on monomeric NLRP3^PYD^ (PDB: 7pdz) and AIM2^PYD^ (PDB: 7k3r), respectively.

### Cell culture and imaging

Each protein was cloned into pCMV6 vector containing C-terminal mCherry (inflammasome PYDs) or eGFP (POPs). HEK293T cells (ATCC, CRL-11268) were seeded into 12-well plate (0.1 x 10^6^ per well) with round cover glass (20 mm). eGFP alone or POP-eGFP plasmids (600 ng and 1200 ng) were co-transfected with inflammasome-mCherry plasmids (300 ng) at 70% confluence using lipofectamine 2000 (Invivogen). After 16 hours, cells were washed twice with 1x phosphate-buffered saline (PBS), fixed with 4% paraformaldehyde, then mounted on glass slides. Images were taken using the Cytation 5 multi-functional reader equipped with a fluorescent microscope (BioTek) and analyzed via the Gen5 software (BioTek). Source Data are appended for each figure.

### Recombinant proteins

Inflammasome proteins were generated and labeled with fluorophores when appropriate as previously described (Matyszewski et al., 2018; Matyszewski et al., 2021; Morrone et al., 2015; Morrone et al., 2014). POP1 was cloned into the pET21b vector, and POP2 and POP3 were cloned into the pET28b vector containing an N-terminal His_6_-MBP tag flanked by a cleavage site for TEVp. All recombinant proteins were expressed in *Escherichia coli* BL21 Rosetta2^DE3^ cells and purified using Ni^2+^-NTA followed by SEC (storage buffer: 40 mM HEPES-NaOH at pH 7.4, 400 mM NaCl, 2 mM dithiothreitol (DTT) 0.5 mM EDTA and 10 % glycerol). Proteins were then concentrated and stored at -80°C.

### Biochemical assays

FRET and FA-based quantitative assays were conducted as described previously (Matyszewski et al., 2018; Matyszewski et al., 2021; Mazanek & Sohn, 2019; Morrone et al., 2014). For example, the polymerization of indicated amounts of FRET-labeled MBP-PYD constructs was triggered by adding TEVp in the presence of increasing amounts of POPs. Half-times for polymerization (t_1/2_s) and the concentration of each POP needed for decreasing the 1/(t_1/2_)s by 50% (IC_50_) was calculated as described in (Matyszewski et al., 2018; Matyszewski et al., 2021). The IC_50_s for inhibiting the dsDNA-binding activities of ALRs were determined with increasing concentrations of POPs as described previously (Matyszewski et al., 2018; Matyszewski et al., 2021); the MBP tag was pre-cleaved in these experiments via TEVp for 30 min. Source Data are appended for each figure.

### nsEM

Each PYD was incubated with TEVp for 30 min to remove MBP tag and promote polymerization in the presence or absence of POPs. Samples were then applied to carbon-coated grids and imaged as described previously (Matyszewski et al., 2018; Matyszewski & Sohn, 2019; Morrone et al., 2015).

## Acknowledgement

We thank Dr. Mariusz Matyzewski and Naveen Mohideen for assistance in Rosetta experiments. Z.M, S.W, J.L, C.M.S, and A.G. conducted *in vitro* and *in cellulo* experiments. G.B., and J.J.Z. performed Rosetta analyses. Z.M., S.W., A.G. interpreted data. J.S. wrote the paper and other authors assisted in editing. This work was supported by NIH R01GM129342 and NSF CAREER (MCB1845003) awards to J.S. Computational Resources were provided by Maryland Advanced Research Computing Center at Johns Hopkins University.

---

**Figure 1 Supplements 1.**
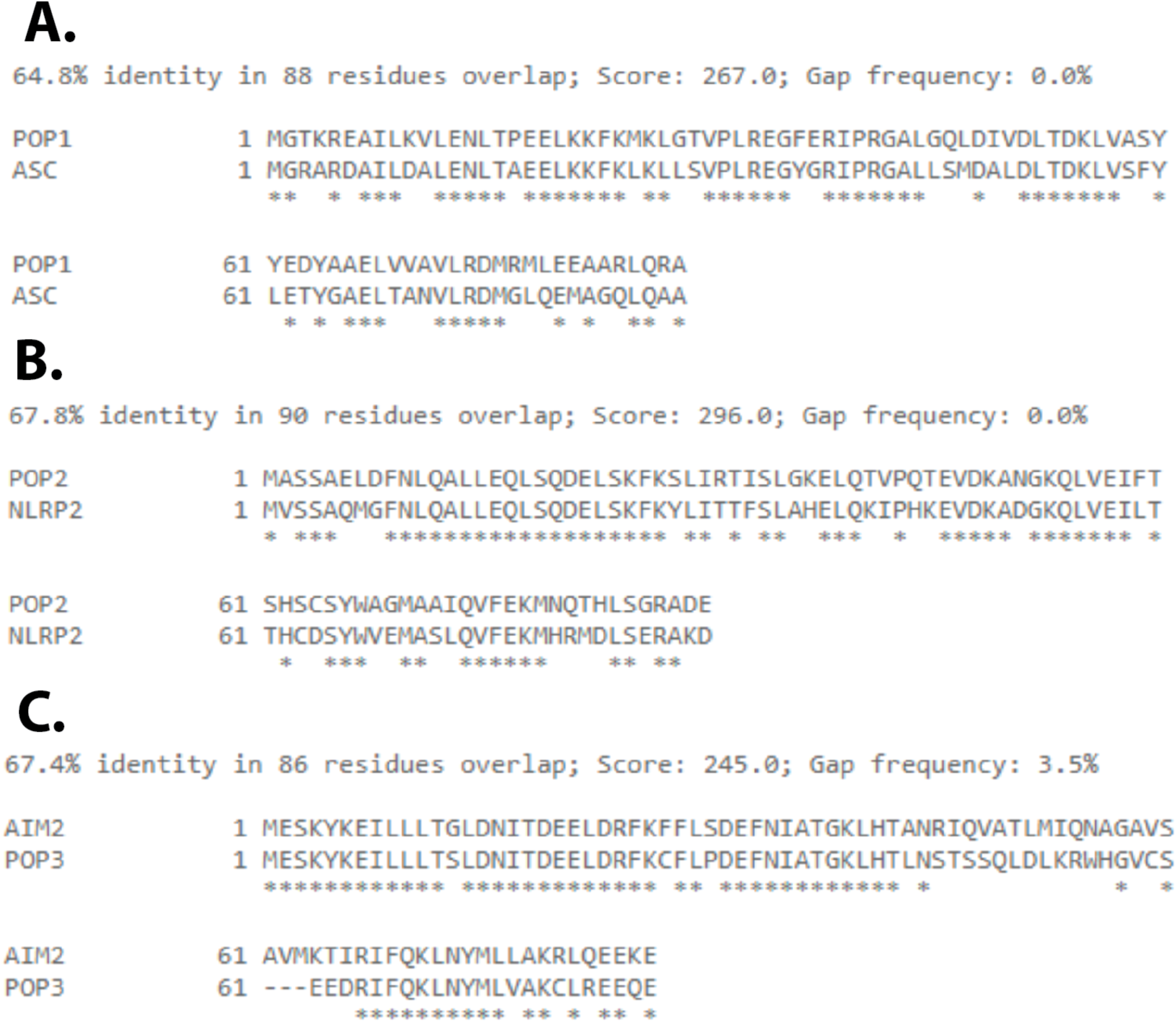
**(A-C)** Amino acid (a.a.) sequence alignments of POPs and their most similar PYDs. See also (de Almeida et al., 2015; Devi et al., 2020; Indramohan et al., 2018; Khare et al., 2014; Ratsimandresy et al., 2017)

**Figure 1 Supplements 2.**
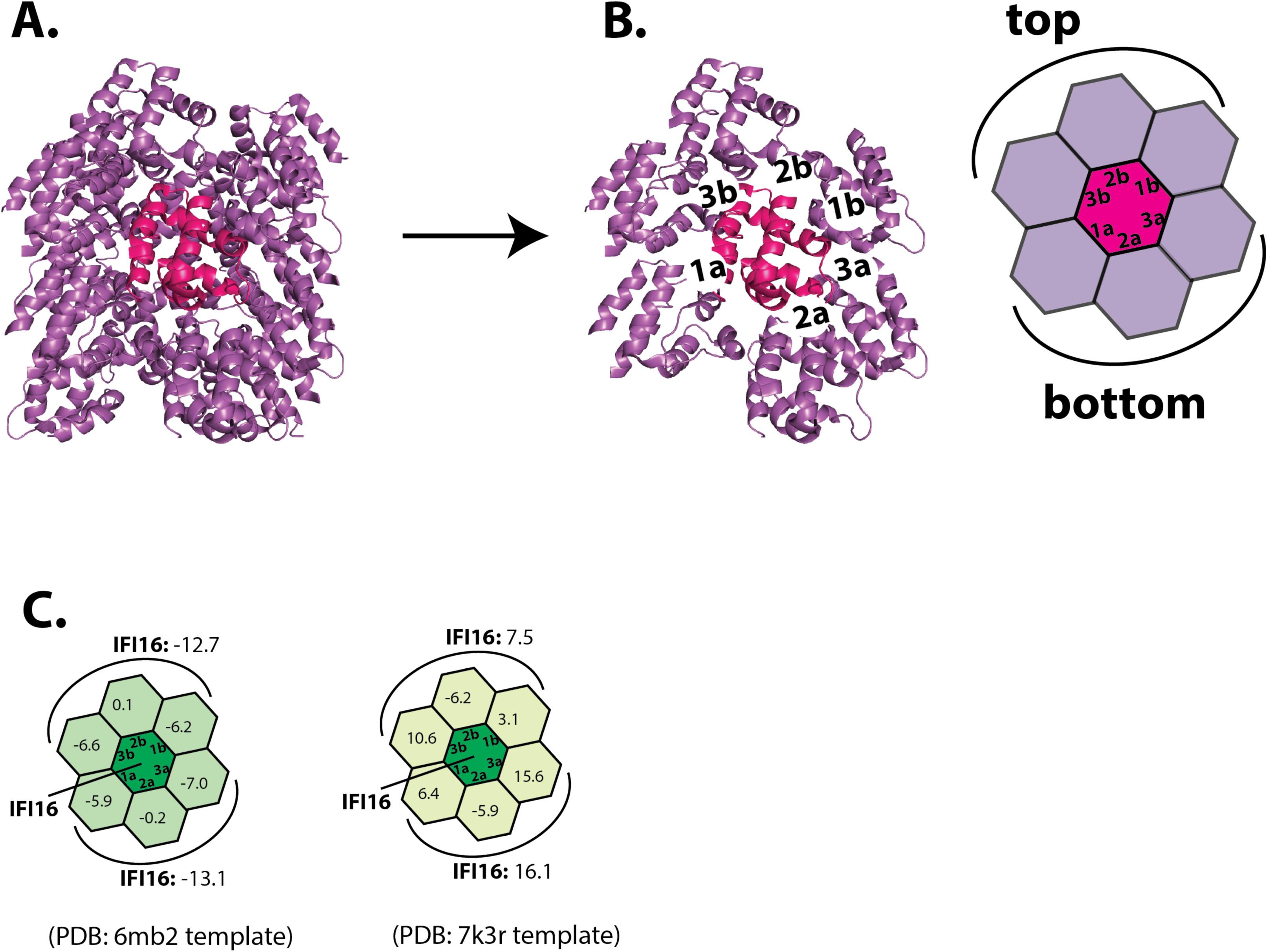
**(A)** The “sideview” of the ASC^PYD^ filament (PDB: 3j63). The center magenta protomer makes all six unique contacts with surrounding purple protomers for assembly. **(B)** The “honeycomb” sideview of the ASC^PYD^ filament (the center protomer and six nearby protomers). Each interface “Type” is labeled, and the corresponding honeycomb diagram is shown on the right. **(C)** IFI16^PYD^ honeycombs based on two different homology models. We used the model based on PDB ID: 6mb2 as it showed more favorable homotypic PYD•PYD interactions (see also Materials and Methods).

**Figure 2 Supplements 1.**
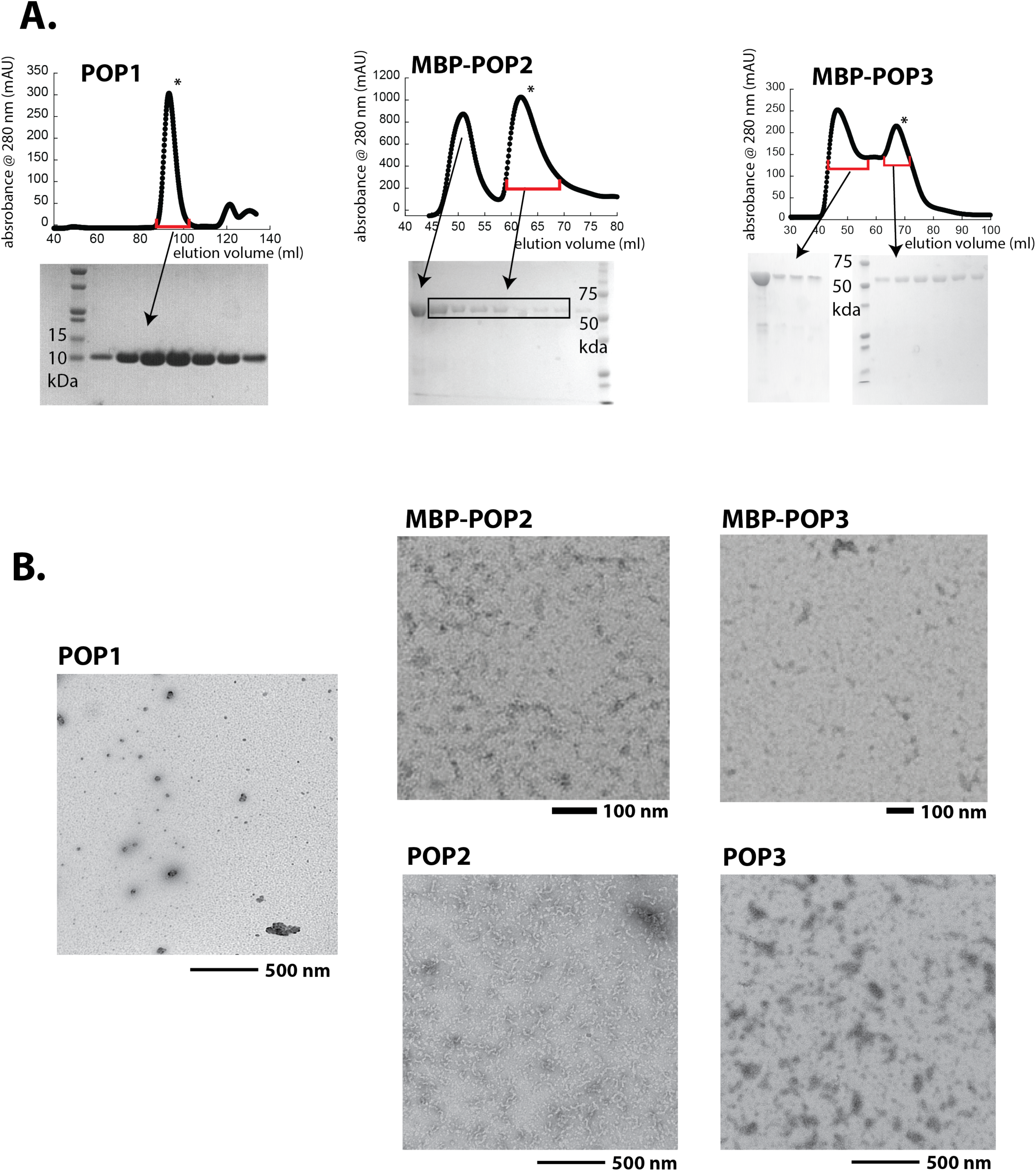
**(A)** Size-exclusion chromatography (SEC; Superdex 75, Cytiva) profiles of recombinant POPs. *: peaks corresponding to monomeric POP1 and MBP-POP2/3 were collected and used in the current study. **(B)** Negative-stain electron microscopy (nsEM) images of untagged POP1 (100 µM), (MBP)- POP2 (20 µM), and (MBP)-POP3 (20 µM). MBP was removed by incubating with TEVp (5 µM) for 30 min.

**Figure 2 Supplements 2.**
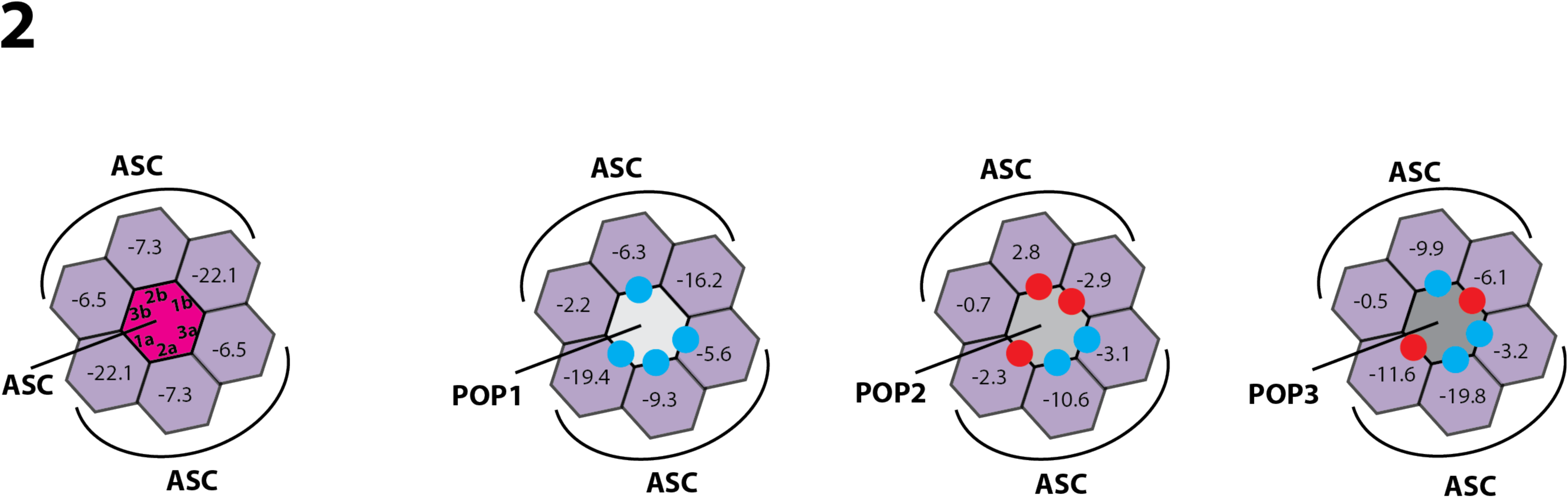
Rosetta interface analysis results showing favorable (ΔΔG ≤ 3.5, blue dots) and unfavorable (ΔΔG ≥ 10.0, red dots) interactions between ASC^PYD^ and POPs.

**Figure 3 Supplements 1.**
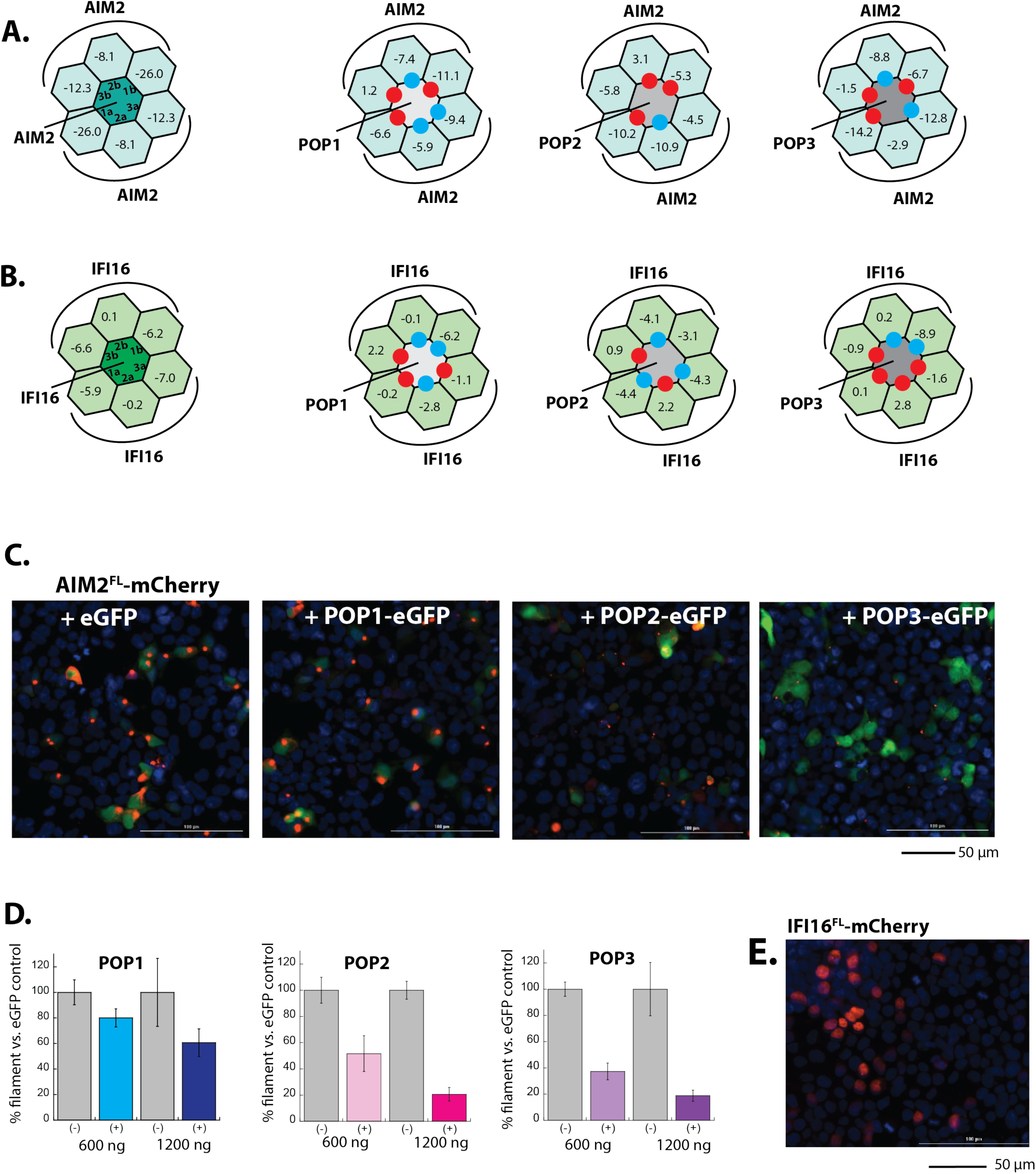
(**A-B**) Rosetta interface analyses results showing favorable (ΔΔG ≤ 3.5, blue dots) and unfavorable (ΔΔG ≥ 10.0, red dots) interactions between AIM2^PYD^ (**A**)/ IFI16^PYD^ (**B**) and POPs. For IFI16, we used ΔΔG ≥ 5.0 or positive ΔGs to identify unfavorable interactions due to the intrinsically weak interface energy scores. **(C)** Sample fluorescent microscope images of HEK293T cells co-transfected with mCherry- tagged AIM2^FL^ (300 ng; crimson) plus eGFP alone or eGFP-tagged POPs (1200 ng; green). Blue: DAPI. **(D)** The relative amounts of AIM2^FL^-mCherry puncta ((300 ng plasmid) in HEK293T cells when co-transfected with POP-eGFP (+) or eGFP alone (-) (600 and 120 ng plasmids). *n* ≥ 4. **(E)** Sample fluorescent microscope images of HEK293T cells transfected with mCherry- tagged IFI16^FL^ (300 ng; crimson). Note the overlap with DAPI staining (blue), thus its nuclear localization.

**Figure 3 Supplements 2.**
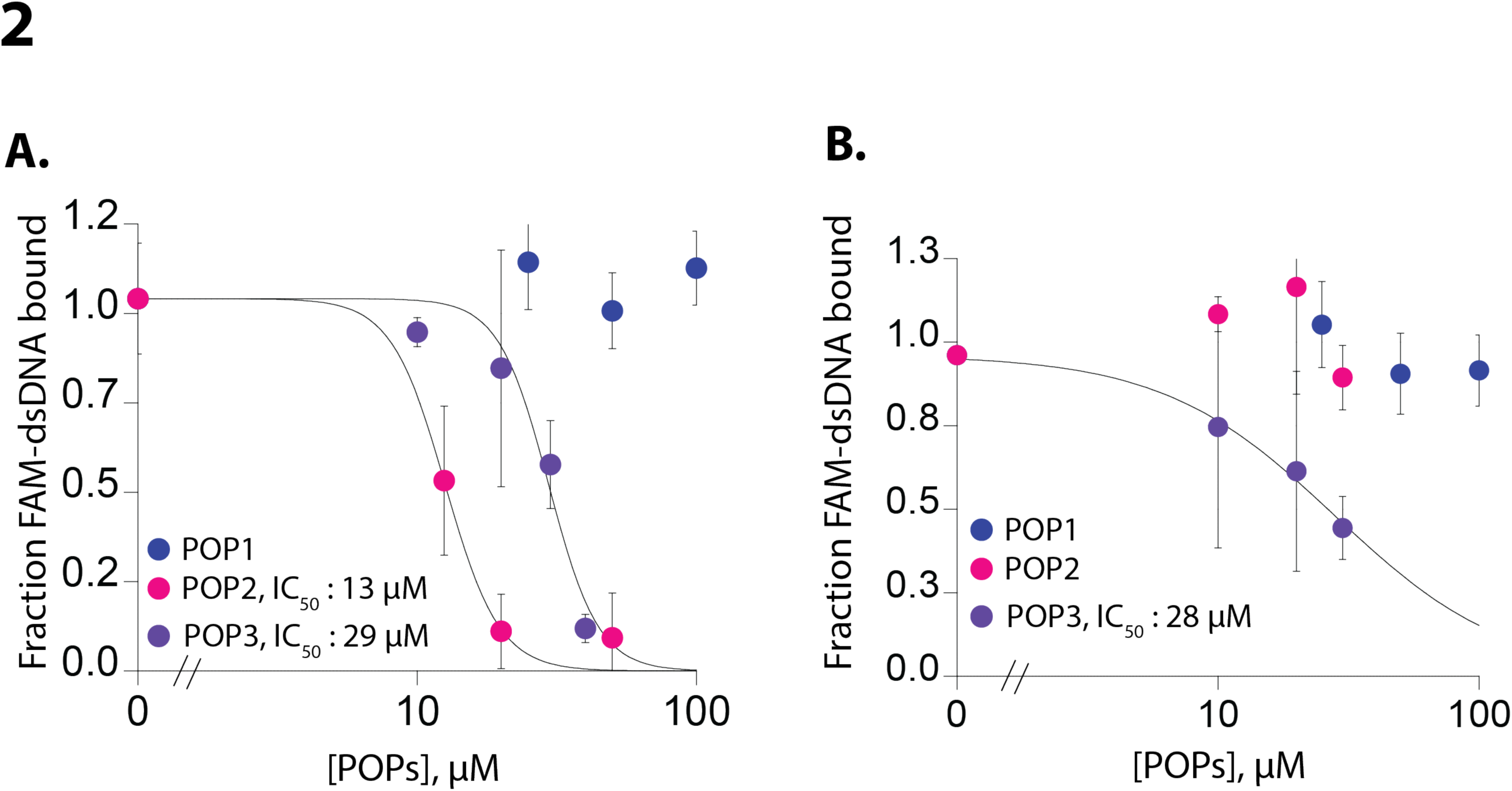
(**A-B**) Plots of fraction fluoresceine-amidate (FAM)-labeled 60-bp dsDNA (10 nM) bound to AIM2^FL^ (100 nM, (**A**)) and IFI16 (200 nM, (**B**)) with increasing concentrations of POPs. Shown is the average of two independent experiments for each POP.

**Figure 4 Supplements 1.**
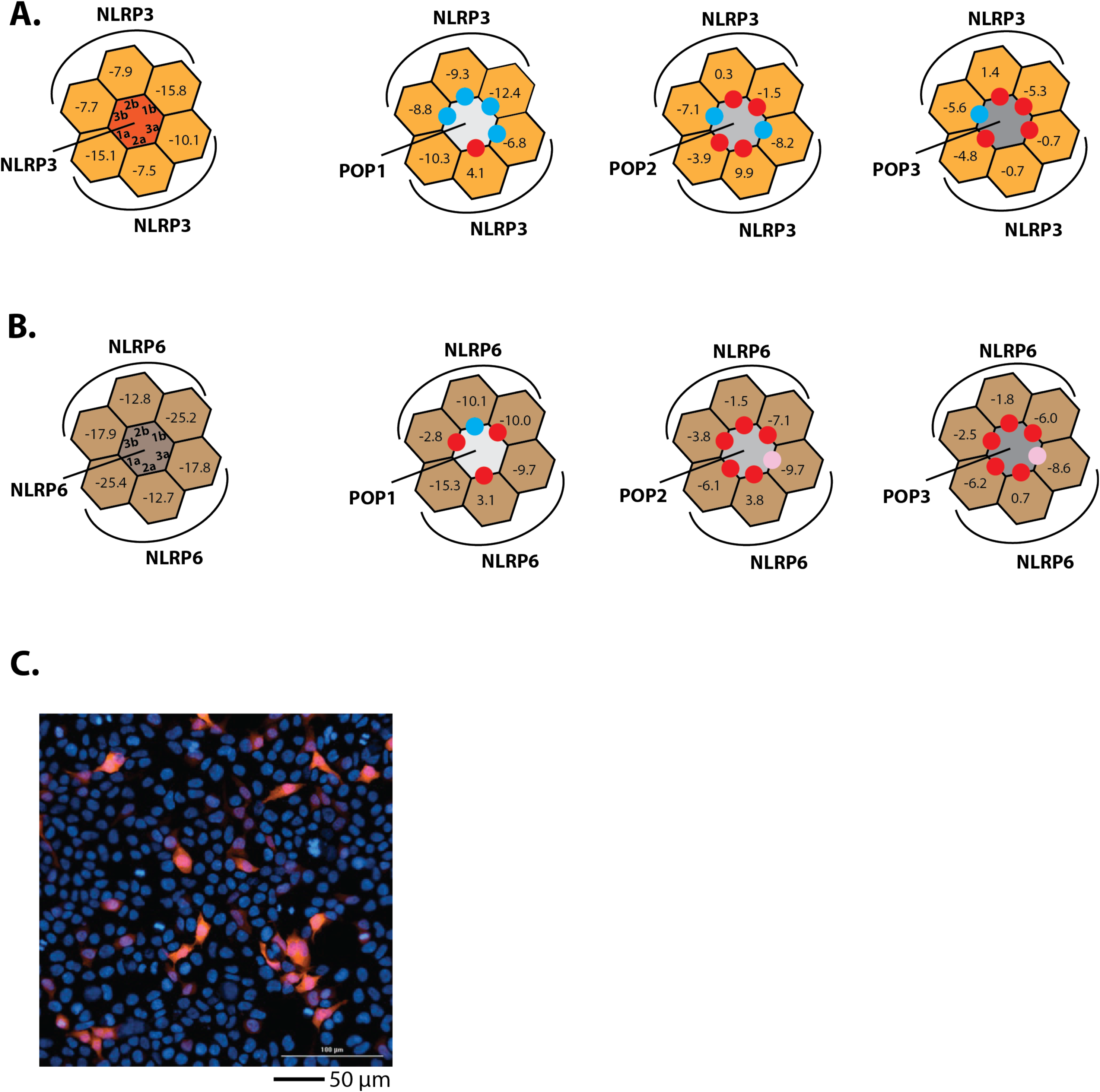
**(A-B)** Rosetta interface analyses results showing favorable (ΔΔG ≤ 3.5, blue dots) and unfavorable (ΔΔG ≥ 10, red dots) interactions between NLRP3^PYD^ (**A**)/ NLRP6^PYD^ (**B**) and POPs. Pink dots indicate interfaces that might allow the recognition between NLRP6^PYD^ and POPs (ΔG ∼9). **(C)** Sample fluorescent microscope images of HEK293T cells co-transfected with mCherry- tagged NLRP2^PYD^ (1000 ng; crimson). Blue: DAPI

**Figure 5 Supplements 1.**
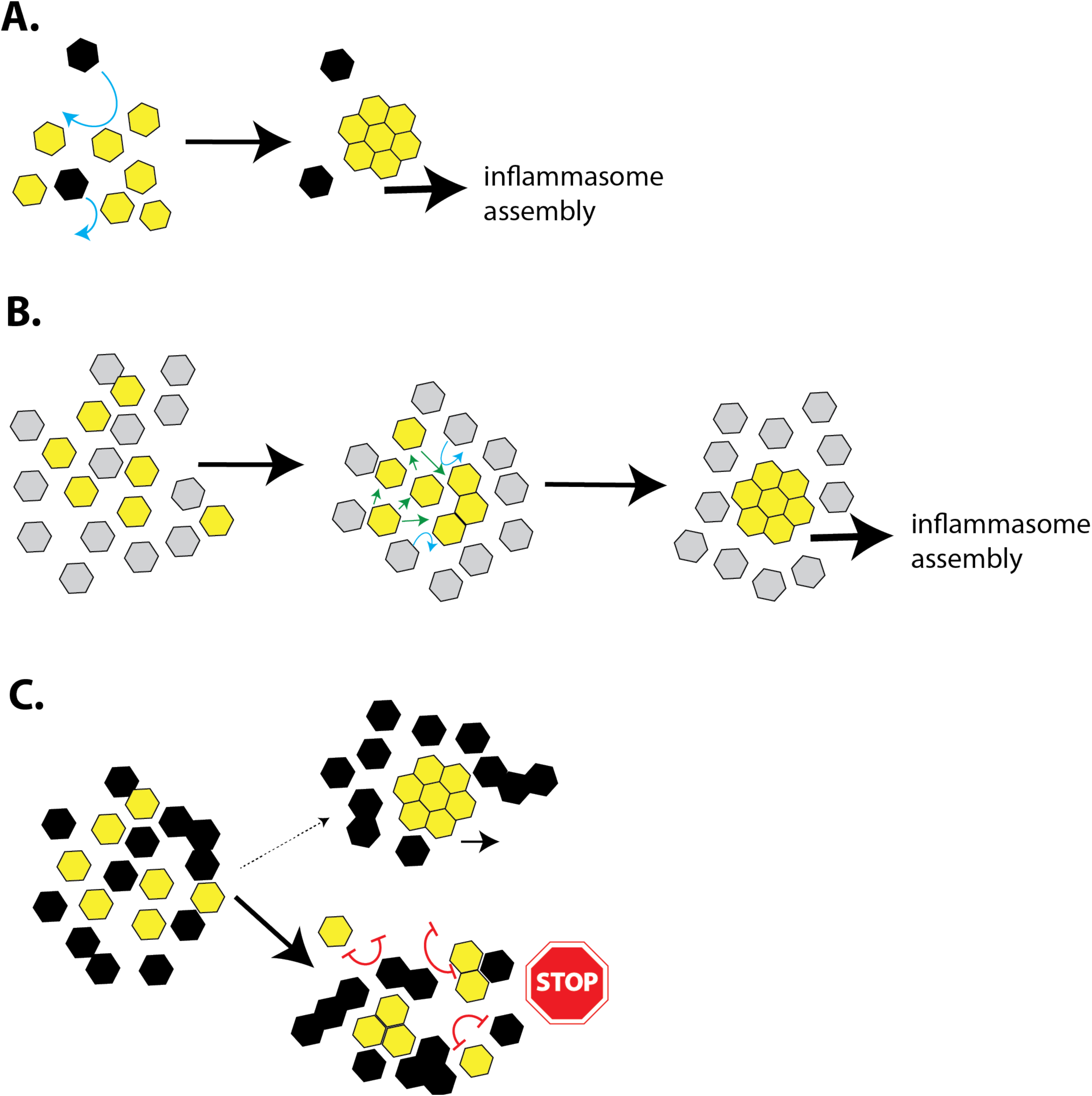
**(A)** A cartoon describing the assembly of inflammasome PYDs in the presence of a basal level of POPs. The more favorable homotypic PYD•PYD interactions would readily outcompete any weak/transient interactions between POPs and PYDs (blue arrows). **(B)** A cartoon describing a possible reason why POP1 cannot inhibit ASC. Because POP1 does not have any strong unfavorable (repulsive) interactions with ASC, the weaker POP•PYD interactions would be easily outcompeted by homotypic PYD•PYD interactions (green arrows) without preventing filament assembly. **(C)** A cartoon describing the inhibition of PYD filament assembly by excess POPs. Favorable ΔGs between POPs and PYDs would allow them to interact transiently, exposing an array of unfavorable (repulsive) interactions that would impede filament assembly (red double- headed blunt arrows). Consequently, inflammasome PYDs would remain either monomeric, or trapped into nonfunctional oligomers. The oligomerization of POP2/3 would also allow multipartite interactions with target PYDs to further enhance their inhibitory effects.

## References

1. Andreeva, L., David, L., Rawson, S., Shen, C., Pasricha, T., Pelegrin, P., & Wu, H. (2021). NLRP3 cages revealed by full-length mouse NLRP3 structure control pathway activation. Cell, 184(26), 6299–6312 e6222. https://doi.org/10.1016/j.cell.2021.11.011

2. Antiochos, B., Matyszewski, M., Sohn, J., Casciola-Rosen, L., & Rosen, A. (2018). IFI16 filament formation in salivary epithelial cells shapes the anti-IFI16 immune response in Sjogren’s syndrome. JCI Insight, 3(18). https://doi.org/10.1172/jci.insight.120179

3. Antiochos, B., Trejo-Zambrano, D., Fenaroli, P., Rosenberg, A., Baer, A., Garg, A., Sohn, J., Li, J., Petri, M., Goldman, D. W., Mecoli, C., Casciola-Rosen, L., & Rosen, A. (2022). The DNA sensors AIM2 and IFI16 are SLE autoantigens that bind neutrophil extracellular traps. Elife, 11. https://doi.org/10.7554/eLife.72103

4. Bae, J. Y., & Park, H. H. (2011). Crystal structure of NALP3 protein pyrin domain (PYD) and its implications in inflammasome assembly. J Biol Chem, 286(45), 39528–39536. https://doi.org/10.1074/jbc.M111.278812

5. Baer, A. N., Petri, M., Sohn, J., Rosen, A., & Casciola-Rosen, L. (2016). Association of Antibodies to Interferon-Inducible Protein-16 With Markers of More Severe Disease in Primary Sjogren’s Syndrome. Arthritis Care Res (Hoboken), 68(2), 254–260. https://doi.org/10.1002/acr.22632

6. Bennett, J. M., Reeves, G., Billman, G. E., & Sturmberg, J. P. (2018). Inflammation-Nature’s Way to Efficiently Respond to All Types of Challenges: Implications for Understanding and Managing “the Epidemic” of Chronic Diseases. Front Med (Lausanne), 5, 316. https://doi.org/10.3389/fmed.2018.00316

7. Brandman, O., & Meyer, T. (2008). Feedback loops shape cellular signals in space and time. Science, 322(5900), 390–395. https://doi.org/10.1126/science.1160617

8. Broz, P., & Dixit, V. M. (2016). Inflammasomes: mechanism of assembly, regulation and signalling. Nat Rev Immunol, 16(7), 407–420. https://doi.org/10.1038/nri.2016.58

9. Cai, X., Chen, J., Xu, H., Liu, S., Jiang, Q. X., Halfmann, R., & Chen, Z. J. (2014). Prion-like polymerization underlies signal transduction in antiviral immune defense and inflammasome activation. Cell, 156(6), 1207–1222. https://doi.org/10.1016/j.cell.2014.01.063

10. de Almeida, L., Khare, S., Misharin, A. V., Patel, R., Ratsimandresy, R. A., Wallin, M. C., Perlman, H., Greaves, D. R., Hoffman, H. M., Dorfleutner, A., & Stehlik, C. (2015). The PYRIN Domain-only Protein POP1 Inhibits Inflammasome Assembly and Ameliorates Inflammatory Disease. Immunity, 43(2), 264–276. https://doi.org/10.1016/j.immuni.2015.07.018

11. Devi, S., Stehlik, C., & Dorfleutner, A. (2020). An Update on CARD Only Proteins (COPs) and PYD Only Proteins (POPs) as Inflammasome Regulators. Int J Mol Sci, 21(18). https://doi.org/10.3390/ijms21186901

12. Fernandes-Alnemri, T., Yu, J. W., Datta, P., Wu, J., & Alnemri, E. S. (2009). AIM2 activates the inflammasome and cell death in response to cytoplasmic DNA. Nature, 458(7237), 509–513. https://doi.org/10.1038/nature07710

13. Franklin, B. S., Bossaller, L., De Nardo, D., Ratter, J. M., Stutz, A., Engels, G., Brenker, C., Nordhoff, M., Mirandola, S. R., Al-Amoudi, A., Mangan, M. S., Zimmer, S., Monks, B. G., Fricke, M., Schmidt, R. E., Espevik, T., Jones, B., Jarnicki, A. G., Hansbro, P. M., … Latz, E. (2014). The adaptor ASC has extracellular and ’prionoid’ activities that propagate inflammation. Nat Immunol, 15(8), 727–737. https://doi.org/10.1038/ni.2913

14. Gong, Q., Robinson, K., Xu, C., Huynh, P. T., Chong, K. H. C., Tan, E. Y. J., Zhang, J., Boo, Z. Z., Teo, D. E. T., Lay, K., Zhang, Y., Lim, J. S. Y., Goh, W. I., Wright, G., Zhong, F. L., Reversade, B., & Wu, B. (2021). Structural basis for distinct inflammasome complex assembly by human NLRP1 and CARD8. Nat Commun, 12(1), 188. https://doi.org/10.1038/s41467-020-20319-5

15. Hochheiser, I. V., Behrmann, H., Hagelueken, G., Rodriguez-Alcazar, J. F., Kopp, A., Latz, E., Behrmann, E., & Geyer, M. (2022). Directionality of PYD filament growth determined by the transition of NLRP3 nucleation seeds to ASC elongation. Sci Adv, 8(19), eabn7583. https://doi.org/10.1126/sciadv.abn7583

16. Hochheiser, I. V., Pilsl, M., Hagelueken, G., Moecking, J., Marleaux, M., Brinkschulte, R., Latz, E., Engel, C., & Geyer, M. (2022). Structure of the NLRP3 decamer bound to the cytokine release inhibitor CRID3. Nature, 604(7904), 184–189. https://doi.org/10.1038/s41586-022-04467-w

17. Hornung, V., Ablasser, A., Charrel-Dennis, M., Bauernfeind, F., Horvath, G., Caffrey, D. R., Latz, E., & Fitzgerald, K. A. (2009). AIM2 recognizes cytosolic dsDNA and forms a caspase-1-activating inflammasome with ASC. Nature, 458(7237), 514–518. https://doi.org/10.1038/nature07725

18. Indramohan, M., Stehlik, C., & Dorfleutner, A. (2018). COPs and POPs Patrol Inflammasome Activation. J Mol Biol, 430(2), 153–173. https://doi.org/10.1016/j.jmb.2017.10.004

19. Iyer, S. S., He, Q., Janczy, J. R., Elliott, E. I., Zhong, Z., Olivier, A. K., Sadler, J. J., Knepper-Adrian, V., Han, R., Qiao, L., Eisenbarth, S. C., Nauseef, W. M., Cassel, S. L., & Sutterwala, F. S. (2013). Mitochondrial cardiolipin is required for Nlrp3 inflammasome activation. Immunity, 39(2), 311–323. https://doi.org/10.1016/j.immuni.2013.08.001

20. Kagan, J. C., Magupalli, V. G., & Wu, H. (2014). SMOCs: supramolecular organizing centres that control innate immunity. Nat Rev Immunol, 14(12), 821–826. https://doi.org/10.1038/nri3757

21. Karki, R., Man, S. M., & Kanneganti, T. D. (2017). Inflammasomes and Cancer. Cancer Immunol Res, 5(2), 94–99. https://doi.org/10.1158/2326-6066.CIR-16-0269

22. Kerur, N., Veettil, M. V., Sharma-Walia, N., Bottero, V., Sadagopan, S., Otageri, P., & Chandran, B. (2011). IFI16 acts as a nuclear pathogen sensor to induce the inflammasome in response to Kaposi Sarcoma-associated herpesvirus infection. Cell Host Microbe, 9(5), 363–375. https://doi.org/10.1016/j.chom.2011.04.008

23. Khare, S., Ratsimandresy, R. A., de Almeida, L., Cuda, C. M., Rellick, S. L., Misharin, A. V., Wallin, M. C., Gangopadhyay, A., Forte, E., Gottwein, E., Perlman, H., Reed, J. C., Greaves, D. R., Dorfleutner, A., & Stehlik, C. (2014). The PYRIN domain-only protein POP3 inhibits ALR inflammasomes and regulates responses to infection with DNA viruses. Nat Immunol, 15(4), 343–353. https://doi.org/10.1038/ni.2829

24. Li, T., Diner, B. A., Chen, J., & Cristea, I. M. (2012). Acetylation modulates cellular distribution and DNA sensing ability of interferon-inducible protein IFI16. Proc Natl Acad Sci U S A, 109(26), 10558–10563. https://doi.org/10.1073/pnas.1203447109

25. Lu, A., Li, Y., Schmidt, F. I., Yin, Q., Chen, S., Fu, T. M., Tong, A. B., Ploegh, H. L., Mao, Y., & Wu, H. (2016). Molecular basis of caspase-1 polymerization and its inhibition by a new capping mechanism. Nat Struct Mol Biol, 23(5), 416–425. https://doi.org/10.1038/nsmb.3199

26. Lu, A., Li, Y., Yin, Q., Ruan, J., Yu, X., Egelman, E., & Wu, H. (2015). Plasticity in PYD assembly revealed by cryo-EM structure of the PYD filament of AIM2. Cell Discov, 1. https://doi.org/10.1038/celldisc.2015.13

27. Lu, A., Magupalli, V. G., Ruan, J., Yin, Q., Atianand, M. K., Vos, M. R., Schroder, G. F., Fitzgerald, K. A., Wu, H., & Egelman, E. H. (2014). Unified polymerization mechanism for the assembly of ASC-dependent inflammasomes. Cell, 156(6), 1193–1206. https://doi.org/10.1016/j.cell.2014.02.008

28. Lu, A., & Wu, H. (2015). Structural mechanisms of inflammasome assembly. FEBS J, 282(3), 435–444. https://doi.org/10.1111/febs.13133

29. Matyszewski, M., Morrone, S. R., & Sohn, J. (2018). Digital signaling network drives the assembly of the AIM2-ASC inflammasome. Proc Natl Acad Sci U S A, 115(9), E1963–E1972. https://doi.org/10.1073/pnas.1712860115

30. Matyszewski, M., & Sohn, J. (2019). Preparation of filamentous proteins for electron microscopy visualization and reconstruction. Methods Enzymol, 625, 167–176. https://doi.org/10.1016/bs.mie.2019.06.007

31. Matyszewski, M., Zheng, W., Lueck, J., Mazanek, Z., Mohideen, N., Lau, A. Y., Egelman, E. H., & Sohn, J. (2021). Distinct axial and lateral interactions within homologous filaments dictate the signaling specificity and order of the AIM2-ASC inflammasome. Nat Commun, 12(1), 2735. https://doi.org/10.1038/s41467-021-23045-8

32. Mayer-Barber, K. D., & Yan, B. (2017). Clash of the Cytokine Titans: counter-regulation of interleukin-1 and type I interferon-mediated inflammatory responses. Cell Mol Immunol, 14(1), 22–35. https://doi.org/10.1038/cmi.2016.25

33. Mazanek, Z., & Sohn, J. (2019). Tracking the polymerization of DNA sensors and inflammasomes using FRET. Methods Enzymol, 625, 87–94. https://doi.org/10.1016/bs.mie.2019.06.006

34. Meizlish, M. L., Franklin, R. A., Zhou, X., & Medzhitov, R. (2021). Tissue Homeostasis and Inflammation. Annu Rev Immunol, 39, 557–581. https://doi.org/10.1146/annurev-immunol-061020-053734

35. Morrone, S. R., Matyszewski, M., Yu, X., Delannoy, M., Egelman, E. H., & Sohn, J. (2015). Assembly-driven activation of the AIM2 foreign-dsDNA sensor provides a polymerization template for downstream ASC. Nat Commun, 6, 7827. https://doi.org/10.1038/ncomms8827

36. Morrone, S. R., Wang, T., Constantoulakis, L. M., Hooy, R. M., Delannoy, M. J., & Sohn, J. (2014). Cooperative assembly of IFI16 filaments on dsDNA provides insights into host defense strategy. Proc Natl Acad Sci U S A, 111(1), E62–71. https://doi.org/10.1073/pnas.1313577111

37. Ohto, U., Kamitsukasa, Y., Ishida, H., Zhang, Z., Murakami, K., Hirama, C., Maekawa, S., & Shimizu, T. (2022). Structural basis for the oligomerization-mediated regulation of NLRP3 inflammasome activation. Proc Natl Acad Sci U S A, 119(11), e2121353119. https://doi.org/10.1073/pnas.2121353119

38. Park, H. H., Logette, E., Raunser, S., Cuenin, S., Walz, T., Tschopp, J., & Wu, H. (2007). Death domain assembly mechanism revealed by crystal structure of the oligomeric PIDDosome core complex. Cell, 128(3), 533–546. https://doi.org/10.1016/j.cell.2007.01.019

39. Periasamy, S., Porter, K. A., Atianand, M. K., H. T. L., Earley, S., Duffy, E. B., Haller, M. C., Chin, H., & Harton, J. A. (2017). Pyrin-only protein 2 limits inflammation but improves protection against bacteria. Nat Commun, 8, 15564. https://doi.org/10.1038/ncomms15564

40. Ratsimandresy, R. A., Chu, L. H., Khare, S., de Almeida, L., Gangopadhyay, A., Indramohan, M., Misharin, A. V., Greaves, D. R., Perlman, H., Dorfleutner, A., & Stehlik, C. (2017). The PYRIN domain-only protein POP2 inhibits inflammasome priming and activation. Nat Commun, 8, 15556. https://doi.org/10.1038/ncomms15556

41. Roberts, T. L., Idris, A., Dunn, J. A., Kelly, G. M., Burnton, C. M., Hodgson, S., Hardy, L. L., Garceau, V., Sweet, M. J., Ross, I. L., Hume, D. A., & Stacey, K. J. (2009). HIN-200 proteins regulate caspase activation in response to foreign cytoplasmic DNA. Science, 323(5917), 1057–1060. https://doi.org/10.1126/science.1169841

42. Sharif, H., Wang, L., Wang, W. L., Magupalli, V. G., Andreeva, L., Qiao, Q., Hauenstein, A. V., Wu, Z., Nunez, G., Mao, Y., & Wu, H. (2019). Structural mechanism for NEK7-licensed activation of NLRP3 inflammasome. Nature, 570(7761), 338–343. https://doi.org/10.1038/s41586-019-1295-z

43. Sharma, M., & de Alba, E. (2021). Structure, Activation and Regulation of NLRP3 and AIM2 Inflammasomes. Int J Mol Sci, 22(2). https://doi.org/10.3390/ijms22020872

44. Shen, C., Li, R., Negro, R., Cheng, J., Vora, S. M., Fu, T. M., Wang, A., He, K., Andreeva, L., Gao, P., Tian, Z., Flavell, R. A., Zhu, S., & Wu, H. (2021). Phase separation drives RNA virus-induced activation of the NLRP6 inflammasome. Cell, 184(23), 5759–5774 e5720. https://doi.org/10.1016/j.cell.2021.09.032

45. Shen, C., Lu, A., Xie, W. J., Ruan, J., Negro, R., Egelman, E. H., Fu, T. M., & Wu, H. (2019). Molecular mechanism for NLRP6 inflammasome assembly and activation. Proc Natl Acad Sci U S A, 116(6), 2052–2057. https://doi.org/10.1073/pnas.1817221116

46. Shi, H., Murray, A., & Beutler, B. (2016). Reconstruction of the Mouse Inflammasome System in HEK293T Cells. Bio Protoc, 6(21). https://doi.org/10.21769/BioProtoc.1986

47. Tartey, S., & Kanneganti, T. D. (2020). Inflammasomes in the pathophysiology of autoinflammatory syndromes. J Leukoc Biol, 107(3), 379–391. https://doi.org/10.1002/JLB.3MIR0919-191R

48. Vora, S. M., Lieberman, J., & Wu, H. (2021). Inflammasome activation at the crux of severe COVID-19. Nat Rev Immunol, 21(11), 694–703. https://doi.org/10.1038/s41577-021-00588-x

49. Zheng, D., Liwinski, T., & Elinav, E. (2020). Inflammasome activation and regulation: toward a better understanding of complex mechanisms. Cell Discov, 6, 36. https://doi.org/10.1038/s41421-020-0167-x

50. Zhong, Z., Zhai, Y., Liang, S., Mori, Y., Han, R., Sutterwala, F. S., & Qiao, L. (2013). TRPM2 links oxidative stress to NLRP3 inflammasome activation. Nat Commun, 4, 1611. https://doi.org/10.1038/ncomms2608

